# Omicron BA.1 breakthrough infection drives long-term remodeling of the memory B cell repertoire in vaccinated individuals

**DOI:** 10.1101/2023.01.27.525575

**Authors:** Aurélien Sokal, Giovanna Barba-Spaeth, Lise Hunault, Ignacio Fernández, Matteo Broketa, Annalisa Meola, Slim Fourati, Imane Azzaoui, Alexis Vandenberghe, Pauline Lagouge-Roussey, Manon Broutin, Anais Roeser, Magali Bouvier-Alias, Etienne Crickx, Laetitia Languille, Morgane Fournier, Marc Michel, Bertrand Godeau, Sébastien Gallien, Giovanna Melica, Yann Nguyen, Florence Canoui-Poitrine, France Noizat-Pirenne, Jérôme Megret, Jean-Michel Pawlotsky, Simon Fillatreau, Claude-Agnès Reynaud, Jean-Claude Weill, Félix A. Rey, Pierre Bruhns, Matthieu Mahévas, Pascal Chappert

## Abstract

How infection by a viral variant showing antigenic drift impacts a preformed mature human memory B cell (MBC) repertoire remains an open question. Here, we studied the MBC response up to 6 months after Omicron BA.1 breakthrough infection in individuals previously vaccinated with three doses of mRNA vaccine. Longitudinal analysis, using single-cell multi-omics and functional analysis of monoclonal antibodies from RBD-specific MBCs, revealed that a BA.1 breakthrough infection mostly recruited pre-existing cross-reactive MBCs with limited *de novo* response against BA.1-restricted epitopes. Reorganization of clonal hierarchy and new rounds of germinal center reaction, however, combined to maintain diversity and induce progressive maturation of the MBC repertoire against common Hu-1 and BA.1, but not BA.5-restricted, SARS-CoV-2 Spike RBD epitopes. Such remodeling was further associated with marked improvement in overall neutralizing breadth and potency. These findings have fundamental implications for the design of future vaccination booster strategies.

## Introduction

Emergence of the SARS-CoV-2 Omicron variant (BA.1) has marked a major antigenic shift in SARS-CoV-2 evolution (van der Straten *et al*, 2022). The Spike (S) protein of SARS-CoV-2 Omicron BA.1 harbors 32 mutations as compared to the ancestral strain (Hu-1) originally identified in Wuhan. These mutations drastically impair neutralizing antibodies elicited by natural infection with the D614G SARS-CoV-2 and/or vaccination with mRNA vaccine encoding the ancestral Hu-1 Spike, and has led to a massive wave of breakthrough infections in the early weeks of 2022 in vaccinated individuals whether they had received 2 or 3 doses of mRNA vaccine (Planas *et al*, 2021; Cameroni *et al*, 2021; Carreño *et al*, 2021; Dejnirattisai *et al*, 2022; Garcia-Beltran *et al*, 2021; Muik *et al*, 2022b). Since, new sub-lineages, displaying additional mutations, continue to emerge supplanting prior variants (Planas *et al*, 2022).

Despite sizeable immune escape by several SARS-CoV-2 variants, the diverse memory B cell (MBC) repertoire generated by two or three doses of mRNA vaccines has been shown to contain high-affinity neutralizing clones against all variants up to BA.1 (Sokal *et al*, 2022; Wang *et al*, 2022a; Goel *et al*, 2022). These MBCs, generated against the ancestral Hu-1 pre-fusion Spike encoded by the original mRNA vaccines, represent an underlying layer of immune protection contributing mostly to the prevention of severe forms of COVID-19 (Dugan *et al*, 2021; Gaebler *et al*, 2021; Rodda *et al*, 2021; Sokal *et al*, 2021b, 2021a). The impact of antigen imprinting in shaping the response and future B cell memory to breakthrough infection by drifted SARS-CoV-2 variants remains an open question of major importance in direct link with the current development of bivalent vaccines and rise in multiple antigenic exposures.

A first hypothesis is that, along re-exposures, the MBCs repertoire will progressively narrow to select mainly broadly reactive B cell receptors (BCR), with limited overall diversity and potential consequences regarding its future ability to cope with new emerging variants. A second hypothesis considers instead that, as previously observed in “primary” infected individuals, exposure to new antigens will engage a de novo naive B cell response with slow maturation through the germinal center, and seeding of new memory B cells into the MBC repertoire, ensuring continued diversity. This latter situation would be in accordance with mouse models showing recruitment of naive B cells in the germinal center during the recall response (Victora & Nussenzweig, 2022). Variant-specific MBCs, targeting mutated residues in the Spike RBD, have been detected in the context of Beta and Gamma SARS-CoV-2 primary infection (Evans *et al*, 2022; Agudelo *et al*, 2022). Recent reports, however, have suggested that the early response occurring in the context of Omicron BA.1 breakthrough infection or Hu-1 mRNA vaccination essentially mobilized cross-reactive clones against conserved Spike glycoprotein epitopes rather than recruiting novel naive B cells specific to mutated BA.1 residues (Kaku *et al*, 2022b; Quandt *et al*, 2022; Muik *et al*, 2022a; Alsoussi *et al*, 2022).

In this study, we longitudinally characterized the humoral response and MBC repertoire after Omicron BA.1 breakthrough infection in a cohort of mRNA vaccinated individuals up to 6 months after infection. We combined single cell multi-omics and functional analysis of several hundred naturally expressed antibodies from RBD-specific MBCs to explore the clonal remodelling, as well as affinity and neutralizing potency evolution of the MBC repertoire. BA.1 breakthrough infection almost exclusively mobilized pre-existing cross-reactive MBCs clones, with limited recruitment of de novo BA.1-restricted responses. Nonetheless, our results demonstrate that reorganization of clonal hierarchy and new rounds of GC reaction combined to maintain diversity and induce progressive maturation of the MBC repertoire against both Hu-1 and BA.1 SARS-CoV-2 Spike RBD variants.

## Results

### Omicron BA.1 breakthrough infection boosts humoral and memory B cell response in triple vaccinated individuals

To understand how the memory B cell repertoire elicited by vaccination is re-shaped by BA.1 breakthrough infection and whether a specific response against its new epitopes occurs, we longitudinally analyzed the SARS-CoV-2-specific B cell responses in 15 individuals with no previous history of COVID-19, which were infected between end of December 2021 and end of January 2022 with Omicron BA.1, shortly after receiving a third dose of BNT162b2 mRNA vaccine (median: 32 days (13-106)). These individuals were sampled at three time points (<1, 2 and 6 months) after BA.1 breakthrough infection to fully characterize the B cell response from the early extra-follicular reaction to the late settlement of long-term memory, combining multiparameter flow cytometry analysis, single-cell RNA-sequencing (scRNA-seq) and single cell culture of spike (S) and RBD-specific B cells (**Figure 1A and Figure S1**). Four of these individuals had been previously sampled after their second and/or third dose of mRNA vaccine (**Table S1)**(Sokal *et al*, 2022, 2023). This provided us with a unique setting to decipher the selection processes occurring at the level of the MBC repertoire upon BA.1 breakthrough infection on a per-individual basis. As control, a parallel cohort of fifteen vaccinated individuals with no history of SARS-CoV-2 infection (SARS-CoV-2-naive) were also sampled at similar time points (<1, 2 and 6 months) after their third dose of mRNA vaccine.

**Figure 1:**
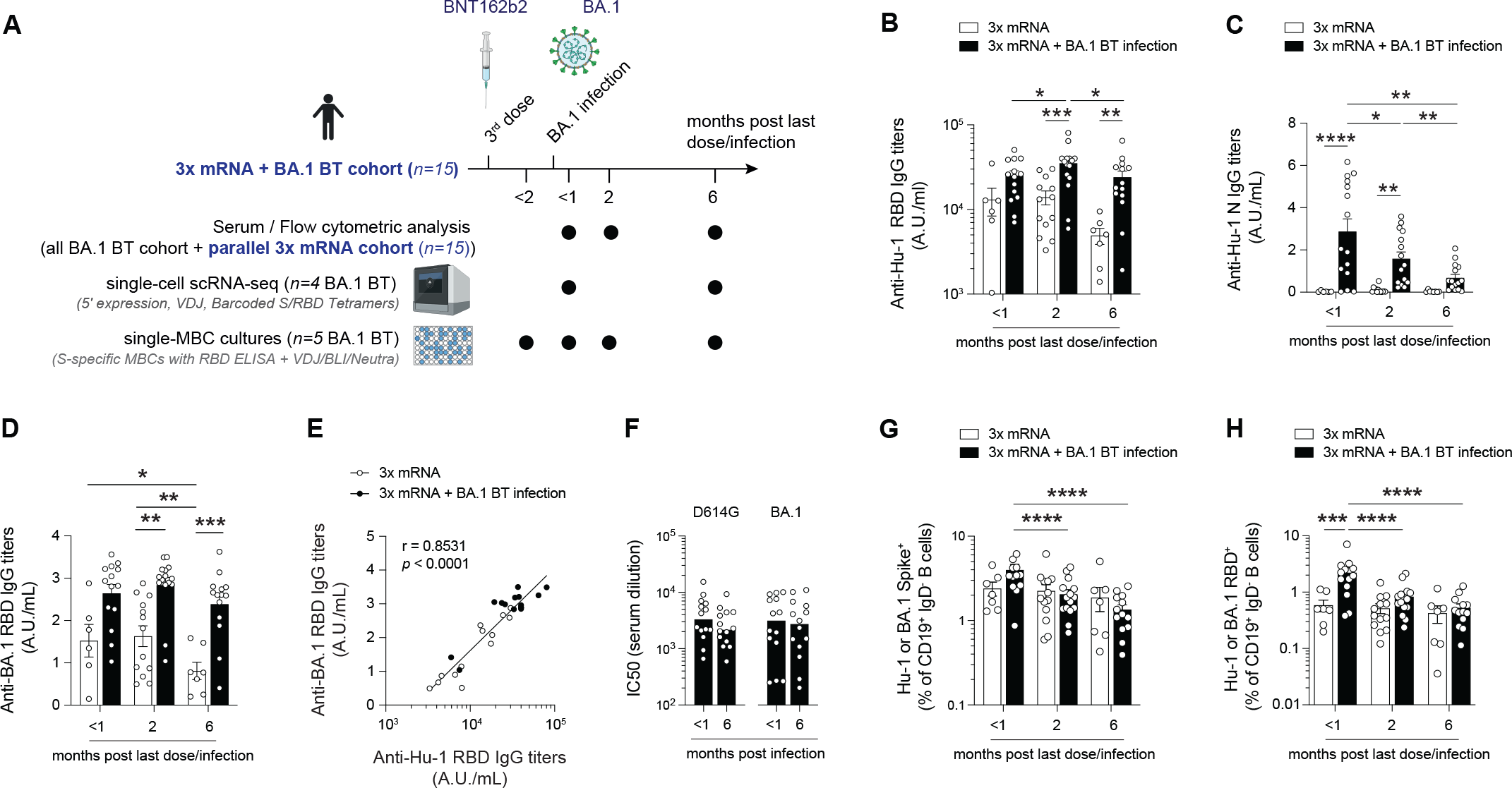
Omicron BA.1 breakthrough infection boosts humoral response in triple vaccinated individuals. (A) Overview of cohort, sampling time points and experimental procedures (see also **Table S1** and **Figure S1A** for detailed sorting strategies). (B, C and D) Anti-SARS-CoV-2 Hu-1 RBD IgG titers (A.U./mL, (B)), anti-Hu-1 Nucleocapsid (N) IgG titers (A.U./mL, (C)) and anti-BA.1 RBD IgG titers (A.U./mL, (D)) in sera from donors after a third dose of mRNA vaccine (white bars, 3x mRNA) or a third dose and subsequent BA.1 breakthrough infection (black bars, 3x mRNA + BA.1 breakthrough (BT) infection) at indicated time-point. Each donor is a dot and bars indicate mean with SEM. (E) Correlation between the anti BA.1 and Hu-1 RBDs IgG titers in sera from 3x mRNA (white dots) or 3x mRNA + BA.1 BT infection donors (black dots) 2 months after last dose/infection. (F) Half maximal inhibitory concentration (IC_50_) for donors’ sera in vitro neutralization assay against authentic D614G (left) or BA.1 (right) SARS-CoV-2 virus at indicated time point after BA.1 BT infection. (G and H) Proportion of all Hu-1 or BA.1 Spike (G) or RBD (H) specific memory B cells among total CD19^+^ IgD^−^ B cells using flow cytometry on PBMCs from 3x mRNA (white bars) or 3x mRNA + BA.1 BT infection donors (black bars) at indicated time-point after last dose/infection (see also **Figure S2A** for detailed gating strategies). In B, C and G and H, we performed mixed model analysis with Tukey’s correction for intra-group comparison and Sidak’s correction for inter-group comparison. In E, non-parametric spearman correlation results on pooled data are represented. In F we performed 2way ANOVA with Sidak’s correction. ****p < 0.0001, ***p < 0.001, **p < 0.01, *p < 0.05. See also **Figure S1**, **Figure S2** and **Table S1**.

Anti-Hu-1 and BA.1 Spike, receptor binding domain (RBD) and Nucleocapsid (N) IgG titers were robustly induced in all individuals after breakthrough infection (**Figure 1B and Figure 1C**), with a good correlation between final anti-Hu-1 and BA.1 RBD titers at the overall cohort level (**Figure 1D and Figure 1E**). N-specific IgG antibodies elicited after BA.1 breakthrough infection waned over the 6 months period, confirming the absence of new SARS-CoV-2 breakthrough in this cohort. The decrease in anti-RBD IgG titers over time was slightly more pronounced in vaccinated SARS-CoV-2-naive individuals, than after BA.1 breakthrough infection, probably reflecting the magnitude of the initial response. The neutralizing activity of plasma against both authentic D614G and BA.1 SARS-CoV-2 strains was high (IC_50_ >100) and, importantly, remained potent against BA.1 at 6 months in all individuals (**Figure 1F**). Longitudinal analyses using a flow panel which included Hu-1 and BA.1 Spike and RBD tetramers, found a major expansion of RBD-specific CD19^+^IgD^−^ B cells shortly after BA.1 breakthrough infection, more pronounced than the Spike-specific response, and with a higher magnitude than that observed after the third mRNA vaccine in SARS-CoV-2-naive individuals (**Figure 1G and 1H**). Both infected and triple vaccinated individuals harbored a sizeable and stable population of Spike and RBD-specific MBCs at the latest time point, after a contraction phase **(Figure 1G and 1H)**. These results show, as previously observed for SARS-CoV-2 (Park *et al*, 2022) or influenza (Wrammert *et al*, 2011), that breakthrough infection in triple vaccinated individuals induces a robust MBC and cross-neutralizing antibody response.

### BA.1 breakthrough infection mobilizes cross-reactive Spike-specific pre-existing MBCs

To characterize the fine specificity and the dynamics of the B cell response after BA.1 breakthrough infection, we first performed multi-parametric FACS analysis on all individuals from both cohorts using major markers of circulating B cell subpopulations (CD19, IgD, CD27, CD38, CD21, CD71 and CD11c) along with Hu-1 and BA.1 Spike and RBD tetramers (**Figures S2A-B**). As previously reported (Kaku *et al*, 2022a; Wang *et al*, 2022b; Quandt *et al*, 2022), BA.1 breakthrough infection mostly mobilized B cells that displayed cross-reactivity against shared epitopes between Hu-1 and BA.1 Spike proteins, representing 70-80% of all RBD-positive cells at any given time point post infection (**Figure 2A and Table S2**). Strikingly, almost no B cells uniquely specific for BA.1 epitopes could be observed both at early time points, as previously described (Kaku *et al*, 2022a; Quandt *et al*, 2022), and at later time points when one can expect to start detecting new germinal center (GC) output (**Figure 2A**). Phenotypic analysis of CD19^+^ IgD^−^ switched B cell populations confirmed a massive expansion of Spike- and RBD-specific CD19^+^IgD^−^CD38^−^CD71^+^ activated B cells (ABCs) occurring in the first couple of weeks post-BA.1 infection (**Figure 2B and 2C; Figures S2C and S2D**), together with the mobilization of CD27^high^ CD38^high^ antibody secreting cells (ASCs) (**Figure S2E**), as previously described upon vaccination or primary infection of SARS-CoV-2 naive individuals (Sokal *et al*, 2021b, 2021a). Activated B cells were enriched in Hu-1/BA.1 cross-reactive cells, confirming the preferential recruitment of these cells in the context of a BA.1 breakthrough infection (**Figure 2D**). Spike- and RBD-specific atypical CD27^−^ IgD^−^ double negative MBCs were also observed, but rarely in CD21^−^ CD11c^−^ DN2 cells (**Figure 2C; Figure S2C**), a population which was previously described as a hallmark of the extra-follicular response in COVID-19 (Woodruff *et al*, 2022, 2020). After their initial expansion, the proportion of Spike- and RBD-specific ABCs decreased over time favoring, in a similar kinetic than observed in triple vaccinated individuals, the resting MBCs, which remain thereafter stable.

**Figure 2:**
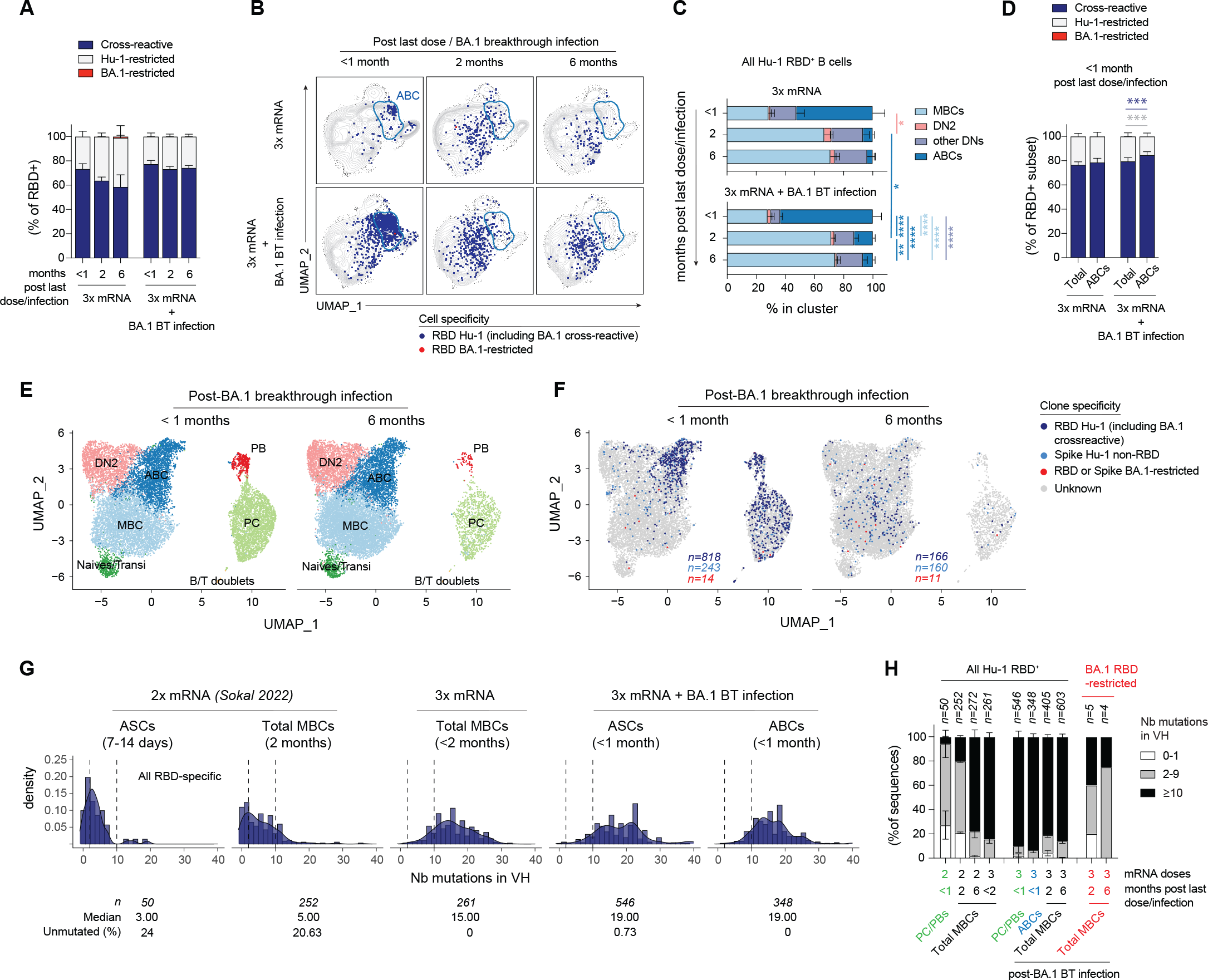
BA.1 breakthrough infection-induced early response recruits Hu-1/BA.1 cross-reactive RBD-specific memory B cells. (A) Frequency of SARS-CoV-2 Hu-1 (Hu-1 only, light grey), Hu-1 and BA-1 (cross reactive, dark blue) or BA.1 (BA.1 only, red) RBD-specific among all RBD-specific CD19^+^ IgD^−^ B cells in donors analyzed by flow cytometry at indicated time point after a third dose of mRNA vaccine (3x mRNA) or a third dose and subsequent BA.1 breakthrough infection (3x mRNA + BA.1 BT infection). (B) UMAP projections of concatenated CD19^+^ IgD^−^ cells from all donors analyzed by multiparametric fluorescence-activated cell sorting (FACS) analysis. RBD-specific B cells are overlaid in dark blue (Hu-1 +/− BA.1 specific) or red (BA.1 only specific) dots on top of all cells from 3x mRNA (top panels) or 3x mRNA + BA.1 BT infection donors (bottom panels) at indicated time point. The ABC cluster is delineated by a blue line. (C) Distribution of RBD specific CD19^+^ IgD^−^ B cells in cluster defined by manual gating strategies (**see Figure S2A**) in 3x mRNA (top panel) and in 3x mRNA + BA.1 BT infection donors (bottom panel) at indicated time point (D) Mean percentage of Hu-1 (Hu-1 only, light grey), Hu-1 and BA-1 (cross reactive, dark blue) or BA.1 (BA.1 only, red) RBD-specific among total (Total) or CD71+ (ABC) RBD-specific CD19^+^ IgD^−^ B cells. (E-F) UMAP of all cells with unique and productive VDJ heavy chain identified in the scRNA-seq analysis of four donors within the 3xmRNA + BA.1 BT infection cohort at the < 1 month (left panel, n=13644) and 6 months’ time points post BA.1 BT infection (right panel, n=14132), with results from unsupervised clustering analysis (E) and cells belonging to clones identified as specific for Hu-1 (+/− BA.1) RBD (dark blue), Hu-1 (+/− BA.1) Spike (S), outside of the RBD, (light blue) or only for BA.1 S or RBD (red) being highlighted (F). (G) Histogram showing the distribution in total number of mutations in the IgV_H_ genes of RBD-specific B cells at indicated time point in indicated population (total MBCs, ABCs or ASCs). Dashed vertical lines indicate 1 and 10 mutations. (H) Bar plots showing the distribution in total number of mutations (0-1: white; 2-9, grey; and ≥10, black) in the IgV_H_ genes of RBD-specific B cells at indicated time point in indicated population. In A, C, D, we performed mixed model analysis with Tukey’s correction for intra-group comparison and Sidak’s correction for inter-group comparison. ****p < 0.0001, ***p < 0.001, **p < 0.01, *p < 0.05. See also **Figure S2 and Table S2 and S3**.

To further get access to the early ASC response, whose heterogeneous surface BCR expression prevents accurate specificity assessment, and to track potential recruitment of naive B cells to the extrafollicular response as well as repertoire and/or transcriptomic changes of the SARS-CoV-2 specific B cell response, we next performed scRNA-seq with parallel VDJ sequencing on sorted CD19^+^IgD^−^ B cells at both early (<1 month) and late time points (6 months) from 4 individuals infected with BA.1 (**Figure S1A**). To focus on cells involved in the ongoing response, CD19^+^IgD^−^ B cells were enriched in Spike- and/or RBD-specific B cells as well as in total antibody-secreting cells (ASCs) (**Figure S1A**). Activated CD19^high^IgD^+^ B cells were also sorted to track potential mobilization of naive B cells (**Figure S1A**). In parallel, we sorted and single-cell cultured Hu-1 and/or BA-1 Spike- and RBD-specific B cells at different timepoints, and IgV_H_ sequences obtained from these cells were further integrated to our scRNA-seq dataset to increase the number of identified Spike- and RBD-specific BCR sequences and add functional information regarding linked antibodies (**Figure S1A-B and Figure S3A**).

Unsupervised clustering analysis of scRNA-seq revealed 6 clusters according to their gene expression profile (**Figure 2E**). Among them, we distinguished CD21^low^CD38^+^CD71^+^ activated B cells (ABCs), CD21^−^CD38^−^CD27^−^CD11c^+^ double-negative 2 (DN2) and 2 clusters of ASCs with both proliferative short-lived plasmablasts (PBs) and non-dividing plasma cells (PCs). The remaining B cells were separated in two populations: a mixture of naive/transitional B cells and a resting MBC population (**Figure 2E**). At early time-point after BA.1 breakthrough infection, Spike- and/or RBD-specific B cells mainly resided among the ABC and ASC clusters (**Figure 2F, Figure S2F, Figure S2G**), and they relocated to the resting MBC cluster at the 6 months’ time point (**Figure 2F**). Concordant with our flow cytometry analysis, these cells were mostly cross-reactive against BA.1 and Hu-1 SARS-CoV-2 with only approximately 1.3 % (14/1,075) of total specific cells analyzed uniquely recognizing BA.1 Spike- or RBD-specific epitopes at any given time point.

Most of the RBD-specific B cells mobilized to the ASC and ABC responses upon BA.1 breakthrough infection harbored a high mutation load (median: 19 mutations), with less than 1% (4/546) unmutated sequences in ASCs and none in ABCs (**Figure 2G and Table S3**) and, overall, very limited frequencies of cells with intermediate level of mutational load (2-9 mutations). Similar results could be observed for S-specific B cells (**Figure S2H-I**). This is in stark contrast with our previous results showing that the RBD-specific ABC and ASC responses after primary infection (Sokal *et al*, 2021b) or 2 dose of mRNA vaccine mobilize cells with low IgV_H_ mutations (**Figure 2G**, Sokal 2022). Non-cross-reactive BA.1 RBD-specific cells appeared to display lower mutational loads (**Figure 2H**), but the very low number of recovered sequences prevented us from drawing any definite conclusion on this point.

Altogether, our results are consistent with a preferential recall of highly mutated pre-existing cross-reactive memory B cells, massively expanding as ABCs and fueling the ASC response, with limited recruitment of naive B cells against novel BA.1 epitopes.

### BA.1 breakthrough infection remodels the MBC repertoire

One of the key questions in the context of immune imprinting relates to understand how a secondary or tertiary antigen encounter reshapes the cognate MBC repertoire and impacts its diversity. First evidence of repertoire remodeling post-BA.1 breakthrough infection could be seen at the global Spike-specific repertoire level, with the proportion of RBD-specific MBCs among Spike-specific clones being significantly increased after BA.1 breakthrough infection (mean±SEM of 51.8±4.2% vs 24.5±3.4%; *p*<0.0001), and remaining significantly higher at 6 months compared in individuals having solely received 3 doses of mRNA vaccine (mean±SEM of 38.8±2.9% vs 20.2±2.4%; *p*=0.0007) (**Figure 3A)**. Further evidence of remodeling could be seen in the RBD-specific MBC repertoire at the clonal level. Longitudinal analysis of the overall RBD-specific MBC clonal diversity, reflected by Chao1 clonal richness index and Shannon entropy values showed no major loss of diversity, apart from the ASCs compartment post-BA.1 infection as expected (**Figure 3B; Figure S3A and S3B**). A sizeable fraction of the RBD-specific clones was maintained over BA.1 infection (“sustained” clones) (**Figures 3C**), most of which being Hu-1/BA.1 cross-reactive (**Figure 3D**). However, a more comprehensive examination of the repertoire revealed marked remodeling at the clonal level, characterized by the loss of previously expanded clones, including Hu-1 RBD-specific only cells, and the emergence of new clones some of which eventually persisted over time (**Figures 3C**). In all longitudinally sampled individuals, sustained clones were still largely represented in the memory B cell repertoire at 6 months (**Figure 3C**), with no clear reduction in the individual frequencies of these clones (**Figure 3E**). Germline V_H_ gene usage in RBD-specific sequences showed no major changes after BA.1 infection relative to those found after a third dose of mRNA vaccine in our longitudinally sampled donors (**Figure 3F**), albeit a progressive enrichment in IGHV1-69 gene usage, as previously described (Kaku *et al*, 2022b), is to be noted.

**Figure 3:**
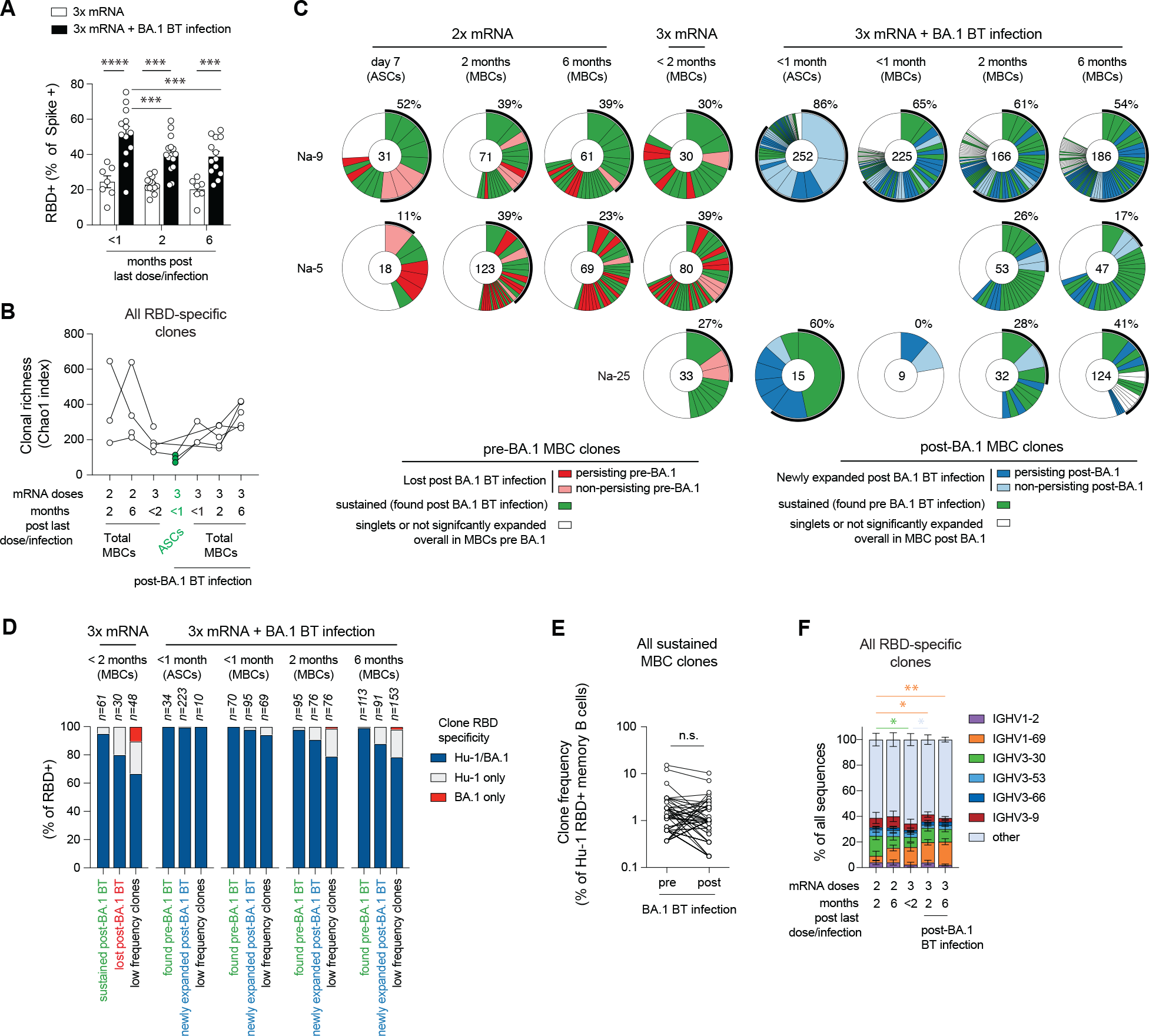
BA.1 breakthrough infection induces partial remodeling of the specific memory B cell repertoire. (A) Frequency of RDB (Hu-1 and/or BA.1) specific CD19^+^ IgD^−^ B cells among Spike (Hu-1 and/or BA.1) specific CD19^+^ IgD^−^ B cells in 3x mRNA (white) or 3x mRNA + BA.1 BT infection (black) cohorts at indicated time points. (B) Evolution over time of clonal richness (Chao1 index) in RBD-specific MBCs (white) or ASCs (green) at indicated time point before and after BA.1 BT. Each line represents one individual donors. (C) Pie charts representing the longitudinal clonal distribution of RBD-specific MBCs and ASC clones in 3 donors from the second dose of mRNA vaccine up to 6 months after BA.1 breakthrough infection. Slice sizes are proportional to the size of each clone. Clones from which members were found before and after BA.1 BT infection are depicted in green. Expanded clones lost upon BA.1 BT infection are represented in light red if found at a single time point or in red if persisting at several timepoint pre-BA.1. Newly expanded clones found after BA.1 BT infection are represented in light blue if found at a single time point or in dark blue if found at several time points. Singletons or expanded clones whose overall frequency post-BA.1 did not reach the frequency of singletons pre-BA.1 are represented in white. Outer black semi-circular line indicates the proportion of sequences belonging to expanded clones at a given time point. The total number of sequences is indicated at the pie center. (D) Frequency of sustained RBD-specific clones among total RBD-specific cells sequenced at any time point pre- or post-BA.1 BT infection. (E) Percentage of cells specific for Hu-1/BA.1, Hu-1 or BA.1 RBD among total RBD-specific cells sequenced, grouped according to their clone’s evolution upon BA.1 BT infection, as defined in (C). (F) IgV_H_ gene usage distribution in CD19^+^IgD^−^ RDB (Hu-1 and/or BA.1) specific B cell after 2 or 3 doses of mRNA vaccine +/− BA.1 BT infection. In A and F, we performed mixed model analysis with Tukey’s correction. In D, we performed a two-tailed Wilcoxon test. ****p < 0.0001, ***p < 0.001, **p < 0.01, *p < 0.05. See also **Figure S3 and Table S3**.

These results suggest that the immune response against the Omicron BA.1 variant does not solely mobilize the top cross-reactive MBCs but also expand memory B cell clones of interest previously found at low frequency in the repertoire or recruits new cross-reactive naive B cells (Figure 3D), maintaining in the process the overall clonal diversity, and thus likely mitigating the negative impact of immune imprinting.

### BA.1 breakthrough infection drives additional affinity maturation and increased overall neutralization breadth of the MBC repertoire

To evaluate the functional consequences of the observed MBC repertoire evolution, we first assayed the affinity against Hu-1, BA.1, BA.2 and BA.5 RBD proteins of over 600 randomly selected monoclonal IgGs, isolated from the supernatants of single cell cultured RBD-specific MBCs from longitudinally sampled donors pre- and post-BA.1 breakthrough infection. Despite being already of high affinity against Hu-1 RBD proteins after 3 doses of mRNA vaccine (median KD of 0.37 ×10^−9^ M), further modest but significant affinity increases of cross-reactive antibodies against Hu-1 and BA.1 RBDs could be observed between 2 and 6 months after BA.1 infection, a trend seen in 4 out of 5 individuals (**Figure 4A; Figure S4A and Table S3**). A similar, albeit non-significant, trend was observed for BA.2 RBD, whereas median affinity of the MBC repertoire rather worsened against BA.5 RBD (Figure 4A). The proportion of MBC-derived mAbs binding both ancestral Hu-1 and BA.1 RBD with similar affinities (“unaffected”, **Figure 4B**) remained stable over time, representing between 60 and 70% of the overall repertoire (**Figure 4C-F**). This suggested that the increase in affinity against BA.1 RBD did not solely result from the selection of clones recognizing unmutated residues. Instead, the overall increase in affinity was nicely reflected both in unaffected and affected clones, with a progressive loss of non-binder/fully impaired clones against the BA.1 RBD (KD>10^−7^ M) between the time point early after the second mRNA vaccine dose and time point late after the BA.1 breakthrough infection (**Figure 4E-F and Figure S4B**). This maturation was not observed against BA.5 RBD, with a small but significant drop in unaffected mAbs at later time points post-BA.1 breakthrough infection (**Figure 4E and Figure S4C**, *p*=0.0196).

**Figure 4:**
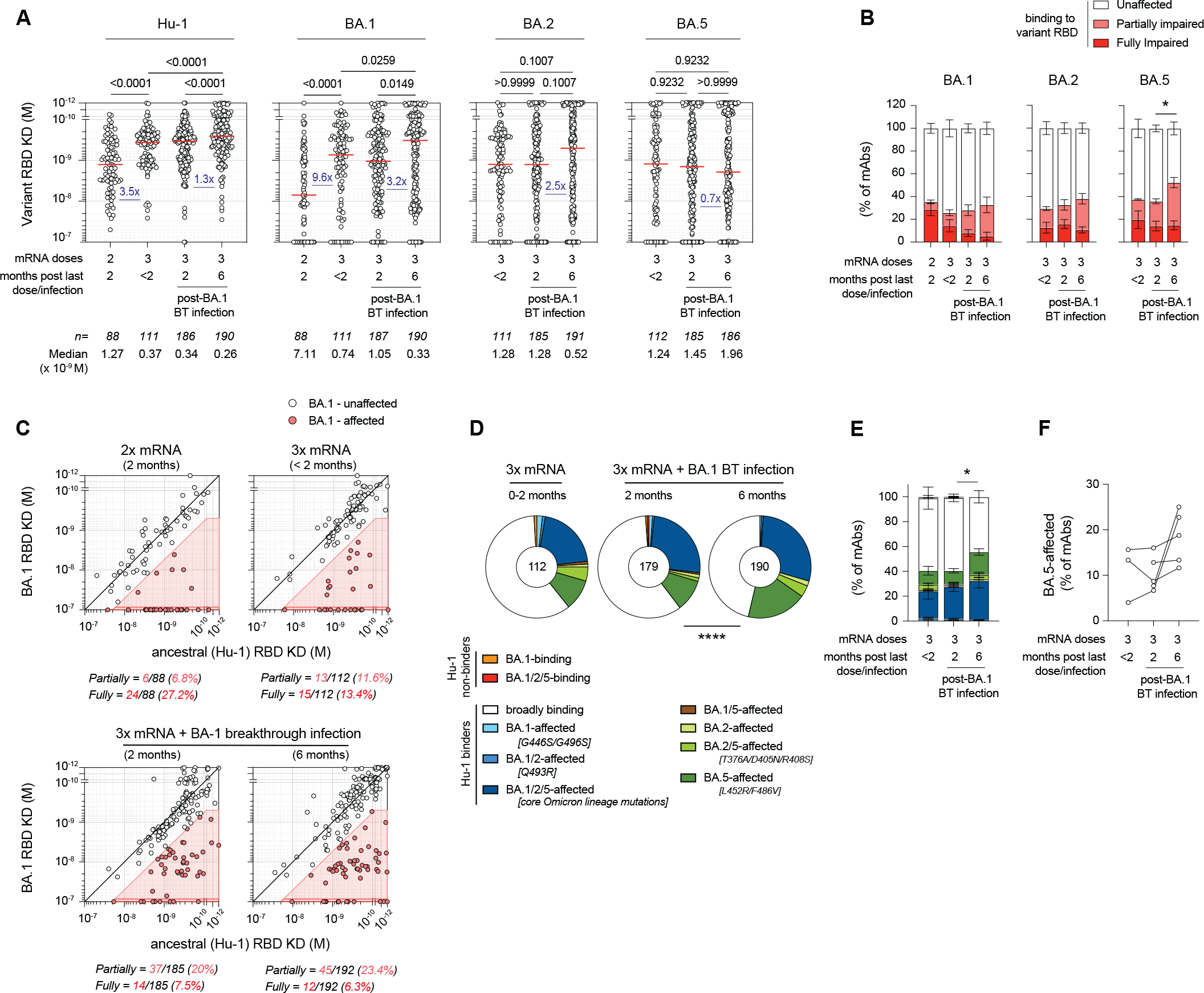
BA.1 breakthrough infection drives additional maturation of Hu-1/BA.1 cross-reactive RBD-specific memory B cells. (A) Dissociation constants (KD, expressed as moles/L) measured by bio-layer interferometry against Hu-1, BA.1, BA.2 and BA.5 RBDs for naturally expressed monoclonal antibodies randomly selected from single-cell culture supernatants of RBD-specific MBCs isolated from 6 donors, including 3 longitudinally analyzed before and after BA.1 BT infection (see **Figure S4**). Numbers of tested monoclonal and median per time point are indicated at the bottom of each graph. (B) Bar plot showing the percentage of tested monoclonals antibodies identified as fully impaired (dark red, variant KD ≥ 10^−7^), partially impaired (ratio of variant/Hu-1 KD > 5 and variant KD ≥ 5×10^−10^, light red) or unaffected (ratio of variant/Hu-1 KD ≤5 or variant KD < 5×10^−10^, white) in their binding to indicated variant RBD. (C) Dot plot representing the KDs for BA.1 versus Hu-1 RBDs for all tested monoclonal antibodies. The red shaded zone indicates BA.1-affected monoclonal antibodies, defined as in (B). (D) Pie charts representing the overall distribution of tested monoclonal according to their binding patterns against Hu-1, BA.1, BA.2 and BA.5 RBDs at indicated time points. Number at the center of the plot indicate the number of tested monoclonal antibodies. (E) Same as in (D) represented as a bar plot and averaged by donor. Bar indicates SEM. (F) Proportion of clones for each donor that are affected in their binding to BA.5 RBD at indicated time point. In A, we performed Kruskal-Wallis tests with Benjamini, Krieger and Yekutieli false discovery rate correction for multiple comparisons (q-values are indicated on the figure). In B and E, we performed mixed model analysis with Tukey’s correction. In D, we performed a Chi-square test. ****p < 0.0001, *p < 0.05. See also **Figure S4 and Table S3**.

Two by two comparisons of binding affinities between Hu-1 and BA.1, BA.2 and BA.5 RBD variants allows to determine the proportion of clones affected by mutations of key binding residues commonly or uniquely shared by Omicron family members (**Figure 4C-E**). We observed few clones affected by BA.1-specific (G446S/G496S), BA.1/BA.2 (Q493R) and BA.2/BA.5-shared (T376A/D405N/R408S) mutations. Most loss or reduction of binding were due to core Omicron lineage (G339D, S375F, K417N, N440K, G446S, S477N, T478K, E484K/Q/A), and BA.5-specific mutations (L452R and F486V), representing approximately 30% and 20% of affected clones at 6 months respectively (**Figure 4D-F**). Interestingly the proportion of monoclonal antibodies solely affected by BA.5 in our analysis (BA-5-affected) appeared to increase between 2 and 6 months. This suggested that the MBC repertoire remodeling occurring after BA.1 breakthrough infection favors clones recognizing conserved immunodominant epitopes in BA.1, which can be mutated in later variants. Germline gene usage according to RBD binding properties did not change over time (**Figure S4D**).

We next evaluated the neutralization potency of these RBD-specific MBCs clones against authentic D614G, BA.1 and BA.5 SARS-CoV-2 viruses. As previously reported, omicron lineage members evaded a large proportion of Hu-1 neutralizing MBCs clones from double or triple vaccinated individuals (**Figure 5A and Table S3**). Neutralization potency of RBD-specific MBCs strongly improved post-BA.1 breakthrough infection against all three strains, with notably 40% of high neutralizing antibodies (>75% at 16nM) and a strong reduction in mAbs with no detectable neutralization potency against BA.1 SARS-CoV-2 (i.e.: <25% neutralization at 16nM) at the latest time point (**Figure 5B)**.

**Figure 5:**
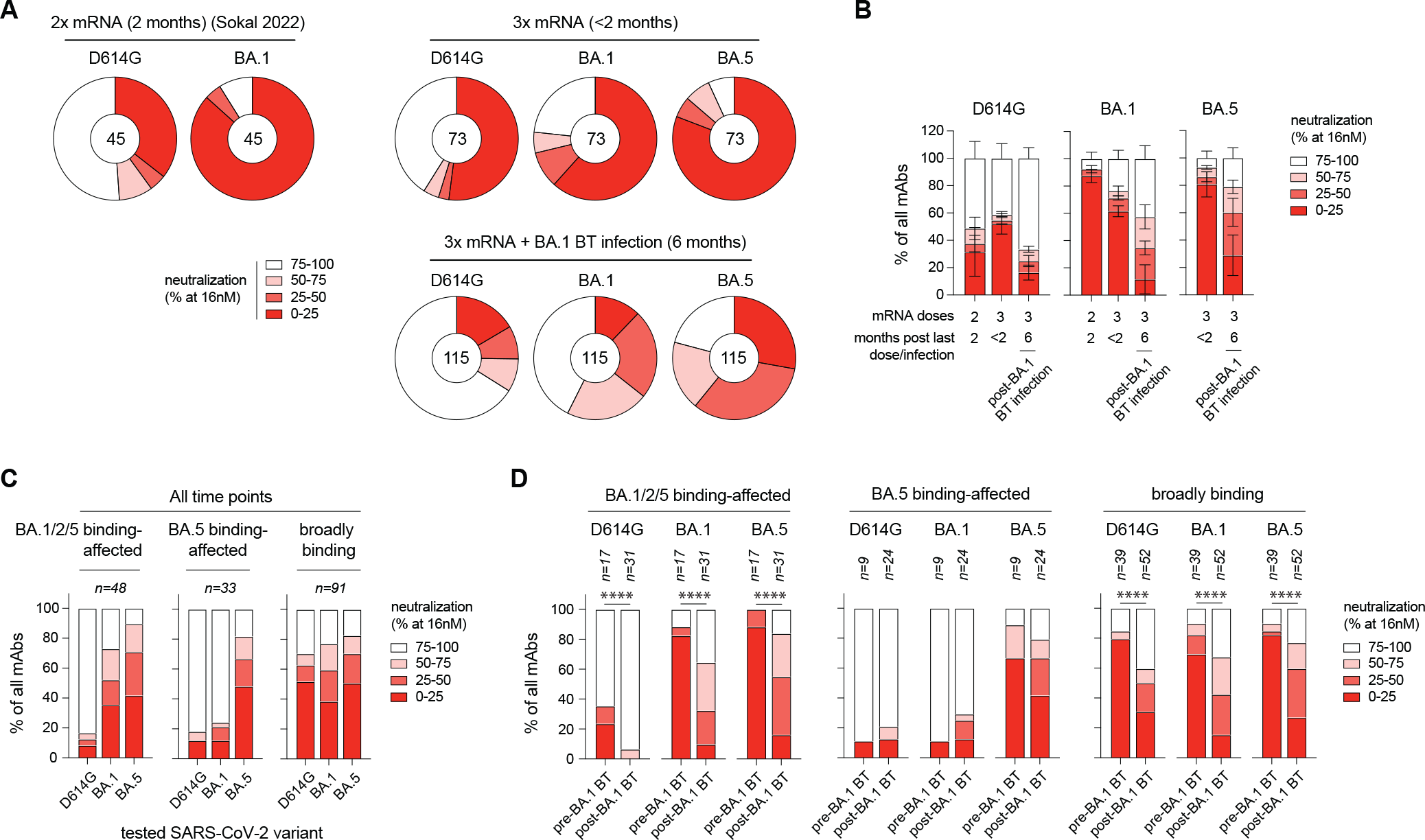
BA.1 breakthrough infection broadly increases neutralization breadth of MBC-derived mAbs. (A) Pie charts representing the distribution in *in vitro* neutralization potency against authentic D614G, BA.1 and BA.5 SARS-CoV-2 virus for naturally expressed monoclonal antibodies at indicated time point randomly selected from RBD specific MBCs, tested at a final concentration of 16 nM. Number at the center of the pie indicates the number of tested monoclonal antibodies. (B) Same as in (A) represented as a bar plot. (C and D) Distribution of the neutralization potency at 16 nM of the tested monoclonal antibodies from MBCs after 3 doses of vaccine, according to their binding pattern to Hu-1, BA.1, BA.2 and BA.5 RBDs (as defined in Figure 4) at all time points (C) or pre and post BA.1 BT (D). In D, we performed Chi-square tests. ****p < 0.0001. See also **Figure S5 and Table S3**.

Cross-examining neutralizing data in the light of RBD affinity measurements showed, as expected, that clones affected in their binding to all tested Omicron RBD variants (BA.1/2/5 binding-affected) lost a large part of their neutralizing potency against both BA.1 and BA.5 variants **(Figure 5C**). Clones affected only in their binding to BA.5 RBD (BA.5 binding-affected), and likely targeting L452R or F486V-containing epitopes, maintained neutralization of the BA.1 strain but lost most of their neutralization potency against the later BA.5 SARS-CoV-2 variant. As previously described (Sokal *et al*, 2022), broadly binding antibodies mostly displayed weak or non-neutralizing potency against all viruses (<50% neutralization at 16nM), suggesting lack of selective pressure. We, however, observed a marked difference between pre- and post-BA.1 tested antibodies in the broadly binding and BA.1/2/5 binding-affected groups (**Figure 5D, Figure S5A**), but not in the BA.5 affected group. This suggests that the observed affinity maturation post-BA.1 breakthrough infection against conserved epitopes as well as common Omicron lineage mutations provide substantial neutralization benefit against future variants sharing similar epitopes.

This substantial gain of neutralization potency was equally observed for neutralizing antibodies using IGHV1-69, IGVH3-30 or IGHV3-53/66 genes, suggesting that these recurrent classes of anti-RBD antibodies can be recruited to participate in neutralization against mutated epitopes (**Figure S5B and S5C)**. Our results thus highlight long-lasting remodeling and affinity maturation of the MBC repertoire following BA.1 breakthrough infection resulting in a clear improvement of the neutralizing overall breadth and potency.

### Selective expansion of MBCs outside GC and additional cycles of GC maturation sequentially contribute to MBC repertoire remodeling

Such remodeling and maturation seen here at the repertoire level post-BA.1 breakthrough infection could be simply explained by the expansion of high-affinity, pre-mutated memory B cells through an extra germinal center reaction (Van Beek *et al*, 2022). Alternatively, this could also reflect a progressive output of clones having undergone new rounds of germinal center reaction, as recently shown in the context of a third mRNA vaccine dose (Alsoussi *et al*, 2022). Increased absolute numbers of total IgV_H_ mutations as well as replacing mutations in the heavy chain complementarity-determining regions (CDRs) of Hu-1/BA.1 cross-reactive RBD-specific ASCs generated after BA.1 infection suggested a preferential recruitment of a highly mutated subpopulation of pre-existing MBCs clones (**Figure 6A and Figure S6A**). In contrast, the longitudinal evolution in mutation profile revealed a decrease in the average number of V_H_ mutations in cross-reactive RBD-specific MBCs at 2 months post-BA.1 infection compared with earlier timepoints, with the presence of 2.77% of unmutated sequences (**Figure 6A**). This drop was also observed at an earlier time point post-infection for replacing mutations in CDRs (**Figure S6A**). This suggested the recruitment of a separate pool of lowly mutated MBCs and/or naive B cells in the MBC pool after BA.1 infection. Subsequently, the average number of IgV_H_ mutations and replacing mutations in CDRs of cross-reactive MBCs significantly increased between 2 and 6 months after BA.1 infection (**Figure 6A, Figure S6A-B**), reaching similar levels as seen before infection.

**Figure 6:**
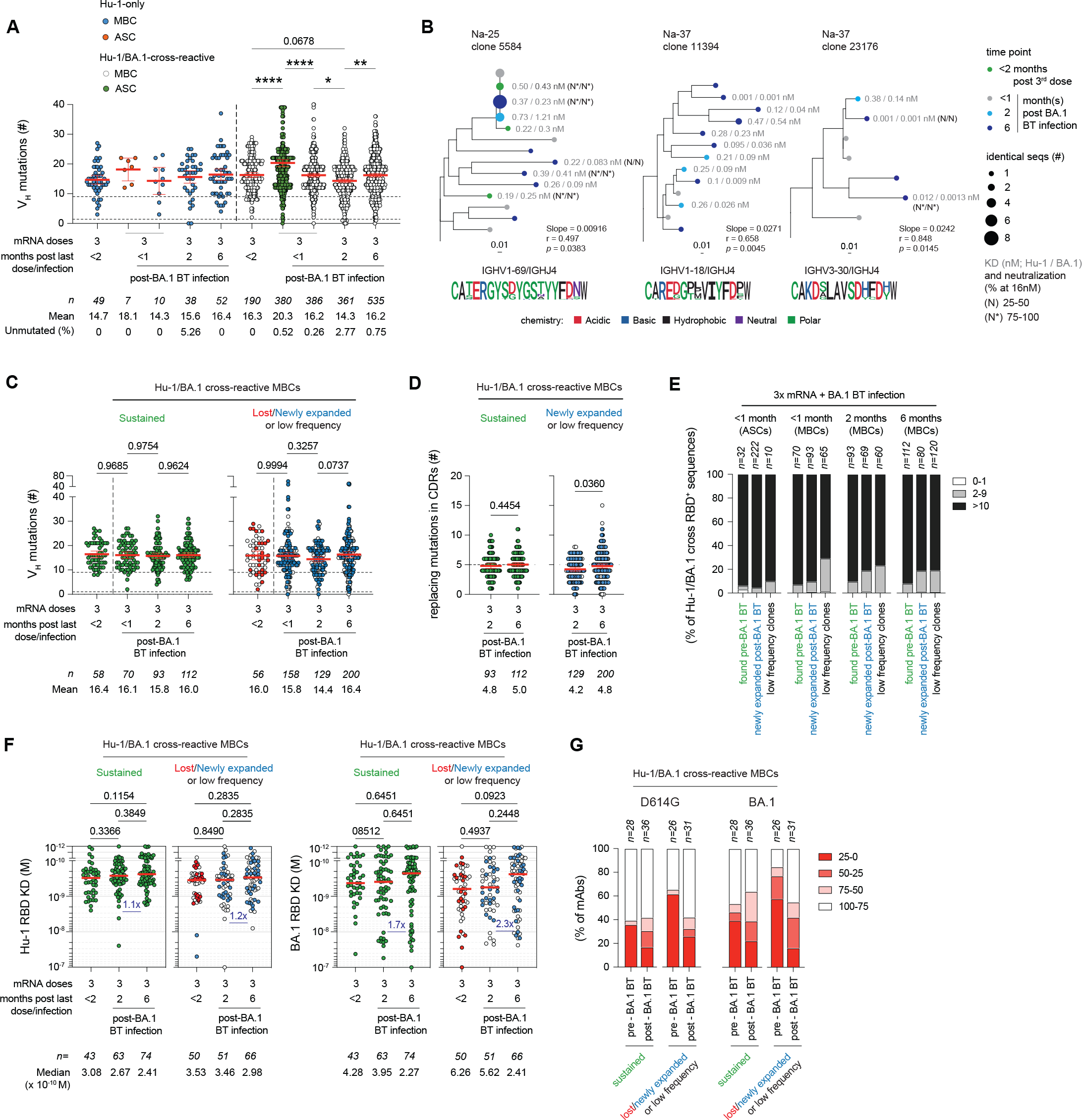
MBC repertoire remodeling and maturation post-BA.1 breakthrough infection reflects successive contributions from extra-GC and GC responses together with the recruitment of low frequency lowly mutated MBC clones. (A) Total number of mutations in the IgV_H_ segment of RBD-specific MBCs or ASCs, sorted according to their specificity for Hu-1 RBD (left panel, blue dots) or cross reactivity (Hu-1 and BA.1 RBDs, white and green dot, right panel), at indicated time points before and after BA.1 BT infection. Mean+/−SEM are shown. (B) Phylogenetic trees for three RBD-specific clones identified as significantly evolving post-BA.1 BT infection, scaled according to the observed frequency of mutations in the IgV_H_ segment. Time point at which the sequences were found are indicated using a color-code and the size of the dot reflect the number of identical sequences found at each time point. At the bottom of each clone, CDR3 (amino acids) from all sequences in the tree are represented as a frequency plot logo. Slope for the rate of somatic mutation accumulation overtime (slope) and p-value (*p*) of the date randomization test comparing the Pearson’s correlation (r) between divergence and time in tree are also indicated. Averaged (median) affinity for Hu-1 (Left) and BA.1 RBDs (Right) are indicated in grey on the right side of each dot corresponding to tested monoclonal antibodies. Neutralization potencies are similarly displayed in black (N: 25-75% and N*: >75% neutralization at 16nM) for both Hu-1 (left) and BA.1 (right) SARS-CoV-2 viruses. (C and D) Number of total (C) or CDR3 replacing mutations (D) in the IgV_H_ segment of Hu-1/BA.1 cross reactive RBD specific MBCs, grouped according to their clone’s evolution upon BA.1 BT infection, as defined in **Figure 3C**, at indicated time point. (E) Bar plots showing the distribution in total number of mutations (0-1: white; 2-9, grey; and ≥10, black) in the IgV_H_ segment of RBD-specific B clones after BA.1 BT infection according to their clone’s evolution upon BA.1 BT infection. (F) Dissociation constants (KD, expressed as moles/L) measured by bio-layer interferometry against Hu-1 (left panels) or BA.1 (right panels) RBDs of Hu-1/BA.1 cross reactive MBCs according to their clone’s evolution upon BA.1 BT infection. (G) In vitro neutralization potency against authentic D614G and BA.1 SARS-CoV-2 virus of naturally expressed monoclonal antibodies from RBD specific MBCs tested at 16 nM and grouped according to sampling time and their clone’s evolution upon BA.1 BT infection. In A and D, we performed ordinary one-way ANOVA with Sidak’s correction for multiple comparisons. In E, we performed unpaired t-tests. In G, we performed Kruskal-Wallis tests with Benjamini, Krieger and Yekutieli false discovery rate correction for multiple comparisons (q-values are indicated on the figure). ****p < 0.0001, **p < 0.01, *p < 0.05. See also **Figure S6 and Table S3**.

This parallel increase in mutational load and affinity maturation observed between 2 and 6 months after BA.1 infection at the level of the MBC repertoire in our longitudinally sampled donors suggested that naive B cells or previously generated MBCs could be recruited to newly formed or vaccine-induced persisting GCs. To investigate this point at the clonal level, we next looked at IgV_H_ mutation number evolution in persisting clones pre- and post-BA.1 infection. Out of 22 total Hu-1/BA.1 cross-reactive RBD-specific clones with more than 7 unique sequences identified at least twice between the post-3rd mRNA vaccine dose time point and later time points after BA.1 breakthrough infection, 3 clones were detected to be significantly accumulating mutations over time (**Figure 6B -Na25-clone 5584**). These numbers, although clearly reduced as compared to the frequency of clones in evolution seen between the 2^nd^ and 3^rd^ vaccine dose (3 out of 4 with more than 7 sequences, **Figure S6C**), are in line with a similar analysis recently performed in the context of influenza vaccine recall response, in which such GC response could be validated by direct staining of draining lymph nodes (Hoehn *et al*, 2021; Turner *et al*, 2020).

One of these three clones identified as evolving (Na-25: clone 5584) also displayed signs of an extra germinal center MBC expansion with 19 cells sharing the exact same sequences that could be detected from the third mRNA vaccine boost to up to 6 months after BA.1 infection. Such identical sequences were observed at an increased frequency of clones in multiple donors at the earlier time points post 2^nd^ and 3^rd^ vaccines doses and post-BA.1 infection (**Figure S6D**), resulting in an overall drop in sequence diversity, with maximal diversity only restored at the 6 months’ time point (**Figure S6E**). This suggests that early and late remodeling in the repertoire post-vaccine boost or post-BA.1 infection likely reflect the combination of an early extra GC response and progressive output from the germinal center.

An additional key hypothesis behind tracking active GC reactions through clonal evolution is that enough sequences of a given clone can be detected at multiple time points. This, however, likely increases the focus on sustained clones (i.e., seen before and after BA.1 infection) at the expense of newly expanded clones that would appear later. Along these lines, sequences with a low level of mutations (<10 mutations) could mostly be seen in low frequency clones early post-BA.1 infection, thereafter slowly transiting to the newly expanded pool (**Figure 6C-E**). This suggest that part of the slight decrease in the overall mutation level in RBD-specific MBCs observed 2 months after infection and the subsequent increase could reflect the recruitment of naive B cells or lowly mutated MBCs previously sparsely represented in the pre-infection repertoire.

At the functional level, however, cells from both sustained and newly expanded/low frequency MBC clones had the tendency to gain in affinity toward the BA.1, but not the ancestral, RBD variant and only 6 months after the breakthrough infection (**Figure 6F**). Nevertheless, we found no systematic correlation between increase in total mutation load and affinity at the repertoire (**Figures S6F-G**) or clonal level, with notably only one out of the three clones identified as statistically increasing their overall mutational load over time after infection also increasing in affinity (**Figure 6B**). Differences, however, were more marked at the level of neutralization potency (**Figure 6G**). While the majority of lost or low-frequency clones pre-BA.1 infection were non or poor-neutralizers of both the D614G and BA.1 SARS-CoV-2 viruses, newly expanded and low-frequency clones seen 6 months after BA.1 infection reached similar neutralization potency as their counterparts from sustained clones.

Overall, these analyses suggest that the MBC repertoire is dynamically reshaped by an early extra-germinal center expansion and subsequent contraction of few selected highly-mutated cross-reactive clones, and the concomitant settlement of a more diverse pool of cells in the repertoire likely, but not necessarily exclusively, representing new GC outputs.

## Discussion

Little is known about the remodeling induced by an infection by a viral variant showing antigenic drift on a repertoire of preformed mature human memory B cells. Selective boosting of cross-reactive antibody specificities by prior exposures was historically coined, in the context of influenza, “original antigenic sin” (Monto *et al*, 2017; Francis, T, 1953). Studies in mice have shown that upon reinfection or re-exposure to an antigen, the MBC pool can expand outside the GCs in an affinity-dependent selective process (Shlomchik, 2018), differentiate in plasma cells, or reenter germinal centers to undergo affinity maturation (Shlomchik & Weisel, 2012; Victora & Nussenzweig, 2022). These secondary germinal centers, however, mostly engage new naive clones allowing diversification against new epitopes (Van Beek *et al*, 2022; Mesin *et al*, 2020; Shlomchik, 2018; Pape *et al*, 2011; Victora & Nussenzweig, 2022). Understanding how these different paths shape the recall response in humans to an antigenic variant of respiratory viruses such as SARS-CoV-2 remains an open question with fundamental implications for the design of future vaccination booster strategies.

As previously reported for influenza (Wrammert *et al*, 2011) and more recently SARS-CoV-2 variant breakthrough infections (Quandt *et al*, 2022; Kaku *et al*, 2022a) or variant-based vaccination (Alsoussi *et al*, 2022), the initial ABC response following BA.1 breakthrough infection is clearly dominated by highly-mutated vaccine-induced cross-reactive MBCs clones eliciting broadly cross-neutralizing antibodies, a point that we could further confirm at the level of ASCs. Here, we show that this imprinting was not limited to the early extrafollicular response, but persisted over time, with very few BA.1-restricted naive B cell clones recruited in de novo germinal centers. High-affinity serum antibodies elicited during the primary response have recently been demonstrated to reduce the recruitment of naive B cells to GCs during secondary responses (Tas *et al*, 2022). Such process, however, is epitope-specific (Tas *et al*, 2022) and, in individuals infected during the first wave of COVID-19, MBCs specific for the spike of seasonal coronaviruses elicited non-neutralizing antibodies against SARS-CoV-2 that did not impair the recruitment of near-germline B cell clones recognizing novel epitopes present in SARS-CoV-2 RBD (Sokal *et al*, 2021; Gaebler *et al*, 2021). Similarly, the massive antibody response against non-neutralizing immunodominant epitopes upon a second immunization with H5N1 vaccine or HIV Env proteins masked these epitopes, thereby promoting maturation of naive B cells in GCs against a different set of non-dominant epitopes (Cirelli *et al*, 2019; Ellebedy *et al*, 2020; Lee *et al*, 2022). In our study, all individuals had recently received a third dose of Hu-1-pre-fusion Spike protein-based mRNA vaccine, but this vaccine boost did not prevent subsequent BA.1 breakthrough infection and MBC recruitment to the extrafollicular response. Omicron’s antigenic distance should thus have enabled exposure of mutated epitopes of the RBD, as previously described in the context of primary infection with SARS-CoV-2 variants (Agudelo *et al*, 2022). One possible explanation for the limited strain-specific response against BA.1 could be the absence of sufficient amount of viral antigen levels to activate naive B cells, as it is rapidly cleared by broadly neutralizing antibodies produced by newly recruited cross-reactive MBCs. Alternatively, the high initial frequency and superior proliferative potential of MBCs may also provide a competitive advantage that restricts antigen accessibility to naive B cells. It remains that we did observe a late recruitment of unmutated and lowly mutated cross-reactive cells in the MBC repertoire. These cells could represent naive B cells recruited to an ongoing GC reaction (de Carvalho *et al*, 2023; Hägglöf *et al*, 2023). This would suggest active selection of cross-reactive B cells in the GC. Alternatively, some of these cells could also be the result of the expansion of pre-existing lowly mutated memory B cells outside any GC reaction.

The absence of de novo recruitment of BA.1-restricted naive B cells and the parallel focus on cross-reactive MBC clones could have induced progressive reduction in overall diversity, a point we only observed transiently upon infection and mostly in the ASCs. And, although non-cross-reactive Hu-1-specific memory B cells tend to be excluded from the early ABC/extra-germinal center response, their frequency returned to pre-BA.1 infection baseline at later time points in the response. Changes in the repertoire up to 6 months after BA.1 infection, however, were not solely restricted to the expansion and later contraction of a cross-reactive MBC response through the extra-germinal center response. The longitudinal tracking of RBD-specific clones revealed a more complex picture with a progressive remodeling of the MBC repertoire, refining clonal hierarchy against BA.1 epitopes, and resulting in a clear improvement in both overall affinity and neutralization breadth. While overall functional variations were clearly more subtle than what can be detected over time in newly vaccinated individuals (Dugan *et al*, 2021; Gaebler *et al*, 2021; Rodda *et al*, 2021; Sokal *et al*, 2021b, 2021a) and this study), our results suggest new cycles of GC maturation for naive B cells and vaccine-induced MBCs following BA.1 breakthrough infection. This is in line with the recent description of a sustained GC response following a vaccine boost in double vaccinated individuals (Alsoussi *et al*, 2022). Additionally, the magnitude of the GC reaction was probably underestimated in our analysis as investigating such marks in the context of a recall response in the PBMCs, with an already fully mature MBC repertoire, is clearly challenging (Hoehn *et al*, 2021). Limited longitudinal sampling of clones of interest, and overrepresentation in the early repertoire post-antigenic exposure of sequences from cells having undergone recent expansion in the extra germinal center response, likely adds up to restrict our analysis to high-frequency persisting clones. The increase in mutational load observed in low-frequency and newly-expanded clones later in the response, clearly suggest a key contribution of these cells to the remodeling of the MBC repertoire. Whether such initially low-frequency clones reenter GCs remains to be demonstrated. This, however, would require direct sampling of draining lymphoid organs. Altogether, the remodeling of the MBC repertoire upon BA.1 breakthrough infection is in line with a recent theoretical modeling study pointing towards the GC reaction during a secondary reaction as a key mechanism to reintroduce diversity in the MBC pool (Van Beek *et al*, 2022). The contribution of the extra-germinal center expansion of MBCs clones and the GCs output to remodel the MBC repertoire appears nevertheless variable from one individual to another.

Finally, our results also raise two clinically relevant points. First, breakthrough infection clearly switched the overall MBC response towards RBD epitopes. Second, post-breakthrough maturation of the MBC response could be considered to expand towards non-mutated immunodominant epitopes on the RBD. The modified pattern of immunodominance at the Spike level could be explained by the epitope masking of highly-conserved region of the S2 domain by the preexisting antibody response (Kaku *et al*, 2022a) or by differential conformational states of SARS-CoV-2 spike protein between the vaccine and the virus. Whether this bears any functional relevance regarding overall protection against future infections, positive or negative, remains to be tested. The modified pattern of immunodominance at the RBD level suggests that repeated challenges with variant Spike proteins may simply increase the selective advantage of specific amino-acid substitutions, as exemplified here for the L452Q/R mutation responsible for part of the immune escape potential of the BA.5 strain from IGHV1-69 and IGHV3-9 class of antibodies (Kaku *et al*, 2022a; Pushparaj *et al*, 2022). In terms of vaccination, it suggests that further strategies to extend the immune response beyond the conserved RBD epitopes will be needed to favor diversity and cope with future antigenic drifts of the SARS-CoV-2.

Collectively these data show that although BA.1 breakthrough infection induce a cross-reactive extra-germinal center expansion, the clonal diversity is maintained by the GCs’ output to remodel and improve neutralization potency and breadth of the MBC repertoire.

## Limitations of the study

Potential limitations of our work include a limited number of subjects included in this study for in depth memory B cell characterization. As such, observed differences between donor groups should be interpreted with caution, notably those groups for which the number of analyzable sequences was low. As previously mentioned, the sparse sampling that can be achieved in peripheral blood may have introduced some bias in clonal representation. Finally, all patients studied were infected with BA.1 early after their third vaccine boost, as has been the case for a sizeable fraction of early Omicron breakthrough infections. In this setting, mRNA vaccine-induced residual germinal centers driven by the Hu-1 pre-fusion Spike were recently activated (Alsoussi *et al*, 2022). It remains to be investigated whether such situation, with two closely related antigens being presented concomitantly as currently implemented in bi-valent vaccines, might favor selection of cross-reactive MBCs.

## Supporting information

Fiugre S4

## Acknowledgement

We thank Garnett Kelsoe (Duke university, Durham, NC, USA) for providing us with the human cell culture system, together with invaluable advice. We thank all the physicians, Constance Guillaud, Raphael Lepeule, Frédéric Schlemmer, Elena Fois, Henri Guillet, Nicolas De Prost, Sarah Feray, Pascal Lim, whose patients were included in this study.

## Author contributions

Conceptualization: P.C., A.S., G.BS, JC.W., F.R., P.B. and M.M.; Data curation: P.C., A.S., L.H., M.B., G.BS, I.A. Formal Analysis: A.S., P.C., L.H., M.B., G.BS, I.A., M.Bo.; A.V., S.F., I.F.; Funding acquisition: S.F., JC.W., CA.R., P.B. and M.M.; Investigation: A.S., P.C., A.DLS., I.A., A.V.; Methodology: P.C., A.S., JC.W., F.R., P.B.,, P.C. and M.M.; Project administration: P.C. and M.M.; Resources: JM.P., F.NP., S.F., E.C., L.G., Ma.Mi., B.G., S.G., G.M., Y.NG., V.Z., P.B., F.R., CA.R., M.M.; Software: P.C; Supervision: P.C., JC.W., P.B., F.R. and M.M.; Validation: A.S., I.A., A.V., M.B., CA.R., PC., MM. ; Visualization: P.C., A.S., I.A., A.DLS., M.M.; Writing – original draft: P.C., A.S., and M.M.; Writing – review & editing: all authors.

---

**Figure S1: Experimental strategy and analysis pipeline for multi-omics-based single cell analysis of SARS-CoV-2-specific B cell response post-BA.1 breakthrough infection. Related to Figure 1**.

(A) Gating strategy used for cell sorting for single cell in vitro culture (left) or single cell RNA sequencing (scRNA-seq, right). For single cell culture, all non-naive B cells (excluding IgD^+^, CD27^−^ naive B cells) binding to the Hu-1 or BA.1 Spike tetramers were single-cell sorted in 96 wells plate pre-coated with CD40L feeders. B cells were defined as live Decoy^−^ CD19^+^ CD3^−^CD14^−^ singlets with additional exclusion of CD38^hi^ cells to remove plasma cells. For scRNA-seq, up to 50.000 live singlets Decoy^−^ IgD^−^CD19^+^CD3^−^CD14^−^ B cells were first sorted. After that, sorting was focused on all remaining Spike-specific (using here four PE-tetramers: Hu-1 and BA.1 RBD and Hu-1 and BA.1 Spike), antibody secreting cells and activated naive B cells (IgD^+^CD19^hi^). (B) Bioinformatic analysis pipeline used for the integration of IgH sequencing and functional data from Spike/RBD -specific MBC single-cell cultures and 10X scRNA-seq VDJ, expression and barcoded surface staining data. **See also Table S1**.

**Figure S2: BA.1 breakthrough infection-induced early response recruits Hu-1/BA.1 cross-reactive Spike-specific memory B cells. Related to Figure 2**.

(A) Gating strategy used for the identification of the main B cell subpopulations as well as SARS-CoV-2 Spike (S) and RBD-specific B cells in our multiparametric FACS analysis. Among CD19^+^IgD^−^ B cells, ASCs were defined as CD19^lo^CD38^hi^ B cells, which could be further subdivided into CD27^−^ early PBs and CD27^+^ PB and PCs. Among non-ASCs cells, activated B cells (ABCs) were first identified as CD71^hi^ and the remaining cells were further subdivided according to their CD27 expression: IgD^−^CD27^+^ resting memory B cells (MBCs) or IgD^−^CD27^−^ double negative B cells (DNs). Among DNs, CD11c^+^CD21^low^ DN2 B cells were further gated. Spike-specific B cells in each subpopulation were defined according to their binding of the Hu-1 and/or BA.1 S tetramers, as shown here in the CD19^+^IgD^−^ B cell pool. Then, among Hu-1 and/or BA.1 S-specific B cell, Hu-1 and/or BA.1 RBD-specific B cells were gated according to their specificity for the Hu-1 and/or BA.1 tetramers. (B) UMAP projections of concatenated CD19^+^IgD^−^ B cells multiparametric fluorescence-activated cell sorting (FACS) analysis. Main B cells subpopulations as defined in (A) are color delineated. (C) Overlay of S-specific (Hu-1 and/or BA.1) B cells on top of multiparametric FACs UMAP.

The ABCs cluster is delineated in blue. (D) Distribution of S-specific CD19^+^IgD^−^ B cells in the cluster defined by the unsupervised FACS analysis in 3x mRNA (top panel) and in 3xmRNA + BA.1 BT infection (bottom panel) at indicated time point. (E) Proportion of Hu-1 and/or BA.1 S-specific cells among ASCs using flow cytometry on PBMCs from 3x mRNA (white bars) or 3x mRNA + BA.1 BT infection donors (black bars) at indicated time-point after last dose/infection. (F) Dot plots showing expression of selected genes in cells from unsupervised scRNAseq clusters. Dot size represents the percentage of cells in the cluster in which transcripts for that gene are detected. Dot color represents the average expression level (scaled normalized counts) of that gene in the population. (G and H) Distribution of all (Hu-1 and/or BA.1) RBD- (G) or S- (H) specific B cells in all identified scRNAseq clusters (top panel) or in MBCs/DN2/ABCs clusters (bottom panel) at indicated time point. (I) Histogram showing the distribution in total number of mutations in the IgV_H_ genes of RBD- and S (outside of the RBD)-specific B cells at the early time point (< 1 month) post-BA.1 BT infection in indicated population (ABCs or ASCs). Dashed vertical lines indicate 1 and 10 mutations. (J) Bar plots showing the distribution in total number of mutations (0-1: white; 2-9, grey; and ≥10, black) in the IgV_H_ genes of RBD-specific B cells at indicated time point in indicated population.

In D and E, we performed mixed model analysis with Tukey’s correction for intra-group comparison and Sidak’s correction for inter-group comparison. ****p < 0.0001, ***p < 0.001, **p < 0.01, *p < 0.05. **See also Table S2 and S3**

**Figure S3: Clonal diversity is maintained over BA.1 BT infection. Related to Figure 3**.

(A) Pie charts representing the clonal distribution of RBD-specific ASCs and MBCs in 6 donors track longitudinally from their second dose of mRNA vaccine and up to 6 months after a BA.1 breakthrough infection. Each slice represents one clone: colored slices indicate expanded clones (2 or more sequences at a given time point) found at several time points in the same individual, gray slices indicate expanded clones found at a single time point, and white slices indicate unique sequences found at several time points. The main white sector in each pie chart represents unique sequences observed at a single time point. The outer black semi-circular line indicates the proportion of sequences belonging to expanded clones at a given time point. The total number of sequences is indicated at the pie center (B) Evolution over time of clonal diversity (Shannon entropy) in RBD-specific MBCs (white) or ASCs (green) at indicated time point before and after BA.1 BT. Each line represents one individual donors. **See also Table S3**.

**Figure S4: BA.1 breakthrough infection drives additional maturation of Hu-1/BA.1 cross-reactive RBD-specific memory B cells. Related to Figure 4**.

(A) Dissociation constants (KD, expressed as moles/L) measured by bio-layer interferometry against Hu-1 and BA.1 RBDs for naturally expressed monoclonal antibodies at indicated time point for each donor analyzed longitudinally. (B) Histograms showing the frequencies of tested monoclonals antibodies that are high-binders (KD < 10^−9^ M), mid-binders (10^−9^ ≤ KD < 10^−8^ M), low-binders (10^−8^ ≤ KD < 10^−7^ M) or considered as non-binders (KD ≥ 10^−7^ M) for BA.1, BA.2 and BA.5 RBDs, at indicated time point. (C) Dot plots representing BA.5 versus Hu-1 RBD KDs for all tested monoclonal antibodies. The red shaded zone indicates BA.5-affected monoclonal antibodies, defined as in **Figure 4B**. (D) Bar plot showing V_H_ gene usage at all timepoints pooled or at indicated time point according to the pattern of binding to BA.1, BA.2 and BA.5 RBDs. Numbers of tested monoclonal antibodies is indicated on top of each bar. **See also Table S3**.

**Figure S5: BA.1 breakthrough infection broadly increases neutralization breadth of MBC-derived mAbs. Related to Figure 5**.

(A) Heatmap representing in vitro neutralization results against D614G, BA.1 and BA.5 SARS-CoV-2 virus for all tested RBD-specific monoclonal antibodies pre- or post-BA.1 BT infection. First annotation line indicates the time point (post-third mRNA dose (< 2 months) or post-third mRNA dose and BA.1 BT infection (6 months). Second annotation line indicates IgV_H_ gene usage. Third annotation line indicates KD for Hu-1 RBD. Fourth annotation line indicates the binding pattern towards omicron lineage RBDs (BA.1, BA.2 and BA.5). The last 3 lines indicate the percent of in vitro neutralization against D614G, BA.1 or BA.5 SARS-CoV-2 as tested at 16nM. (B) In vitro neutralization potency at 16 nM against D614G, BA.1 and BA.5 SARS-CoV-2 virus according to IgV_H_ usage for all tested monoclonal antibodies from RBD-specific MBCs (pooled from all time points). (C) In vitro neutralization potency at 16 nM against D614G, BA.1 and BA.5 SARS-CoV-2 for monoclonal antibodies using selected IgV_H_, further grouped based on time of sampling (before or after the BA.1 BT infection). **See also Table S3**.

**Figure S6: MBC repertoire remodeling and maturation post-BA.1 breakthrough infection reflects successive contributions from extra-GC and GC responses together with the recruitment of low frequency lowly mutated MBC clones. Related to Figure 6**.

(A) Violin plots showing total numbers of replacing mutations in IgV_H_ CDRs regions of Hu-1 and BA.1 RBD-specific (cross-reactive) MBCs (white) or ASCs (green) at successive time points before and after BA.1 BT infection. Red line indicates mean +/− SEM. (B) Violin plots showing total numbers of mutations in the IgV_H_ segment of BA.1 and Hu-1 RBD-specific (cross reactive) MBCs at successive time points before and after BA.1 BT infection in individual donors. Red line indicates mean +/− SEM. (C) Phylogenetic trees for three RBD-specific clones identified as significantly evolving post-second mRNA vaccine dose and pre-BA.1 BT infection, scaled according to the observed frequency of mutations in the IgV_H_ segment. Time point at which the sequences were found are indicated using a color-code and the size of the dot reflect the number of identical sequences found at each time point. At the bottom of each clone, CDR3 (amino acids) from all sequences in the tree are represented as a frequency plot logo. Slope for the rate of somatic mutation accumulation overtime (slope) and p-value (*p*) of the date randomization test comparing the Pearson’s correlation (r) between divergence and time in tree are also indicated. Averaged (median) affinity for Hu-1 (Left) and BA.1 RBDs (Right) are indicated in grey on the right side of each dot corresponding to tested monoclonal antibodies. Neutralization potencies are similarly displayed in black (N: 25-75% and N*: >75% neutralization at 16nM) for both Hu-1 (left) and BA.1 (right) SARS-CoV-2 viruses. (D) Estimated frequency of identical IgV_H_ sequences found in RBD-specific MBCs (white) or ASCs (green) at indicated time point before and after BA.1 BT infection. Estimated frequency correspond to the average frequency of identical sequences found upon 1000 bootstrapping, downsampling to 65 sequences per time point for each donor to correct for sampling biases. (E) Evolution over time of sequence richness (Chao1 index) in RBD-specific MBCs (white) or ASCs (green) at indicated time point before and after BA.1 BT. Each line represents one individual donors. (F and G) Plot showing measured Hu-1 (F) or BA.1 (G) RBD KDs versus the number of mutations identified in the IgV_H_ segment of RBD-specific MBCs in sustained (left panels) or lost/newly expanded or low frequency clones (right panels). Dot color indicates the time point from which each monoclonal antibody was sampled (red: 2 months after second mRNA dose, white: <2 months after third mRNA dose, light blue: 2 months after BA.1 BT infection, dark blue: 6 months after BA.1 BT infection).

In A and B, we performed ordinary one-way ANOVA with Sidak’s correction for multiple comparisons. In E, we performed paired t-tests. ****p < 0.0001, ***p < 0.001, **p < 0.01, *p < 0.05. **See also Table S3**.

## STARMethods

### Key Resource Table

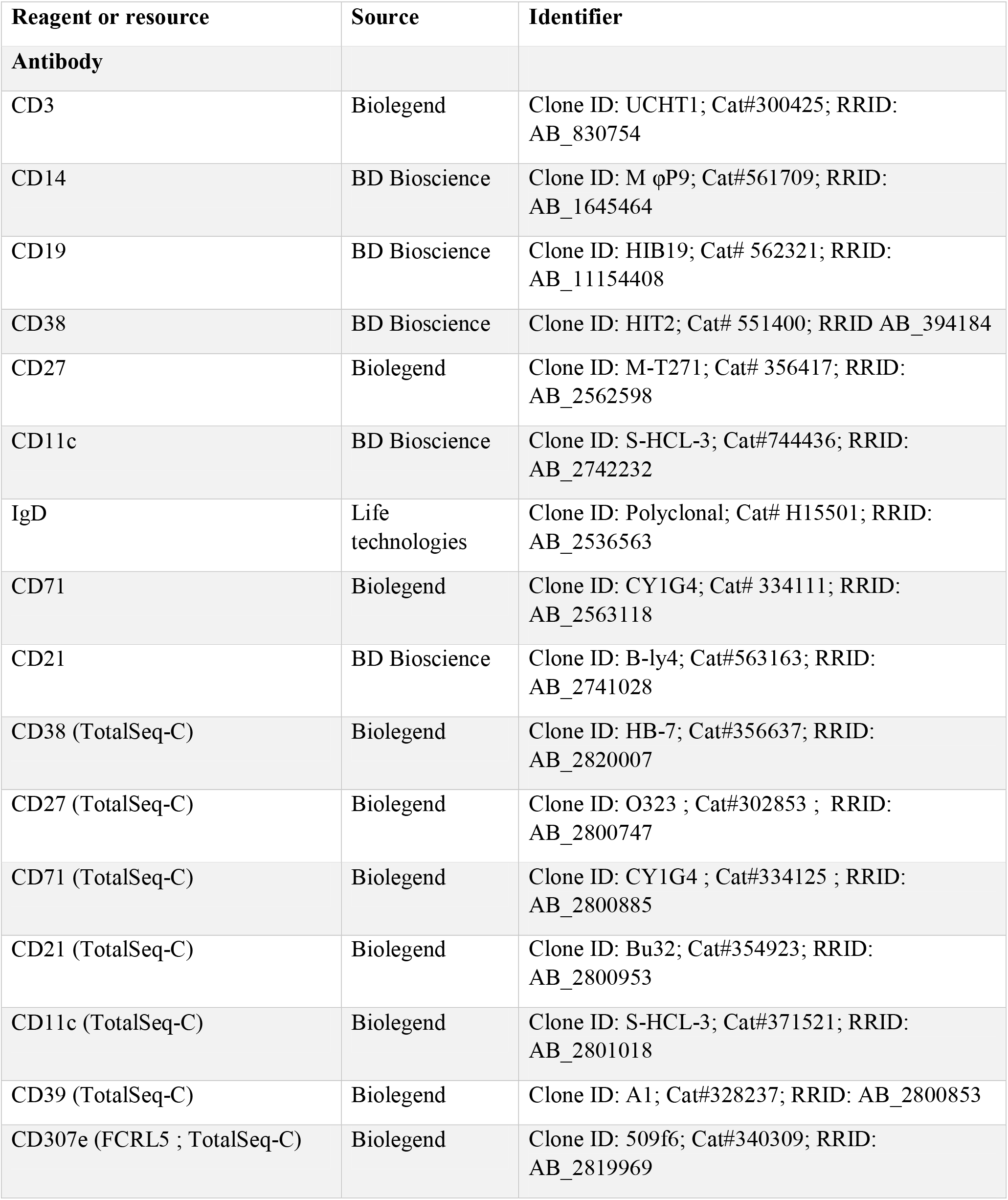
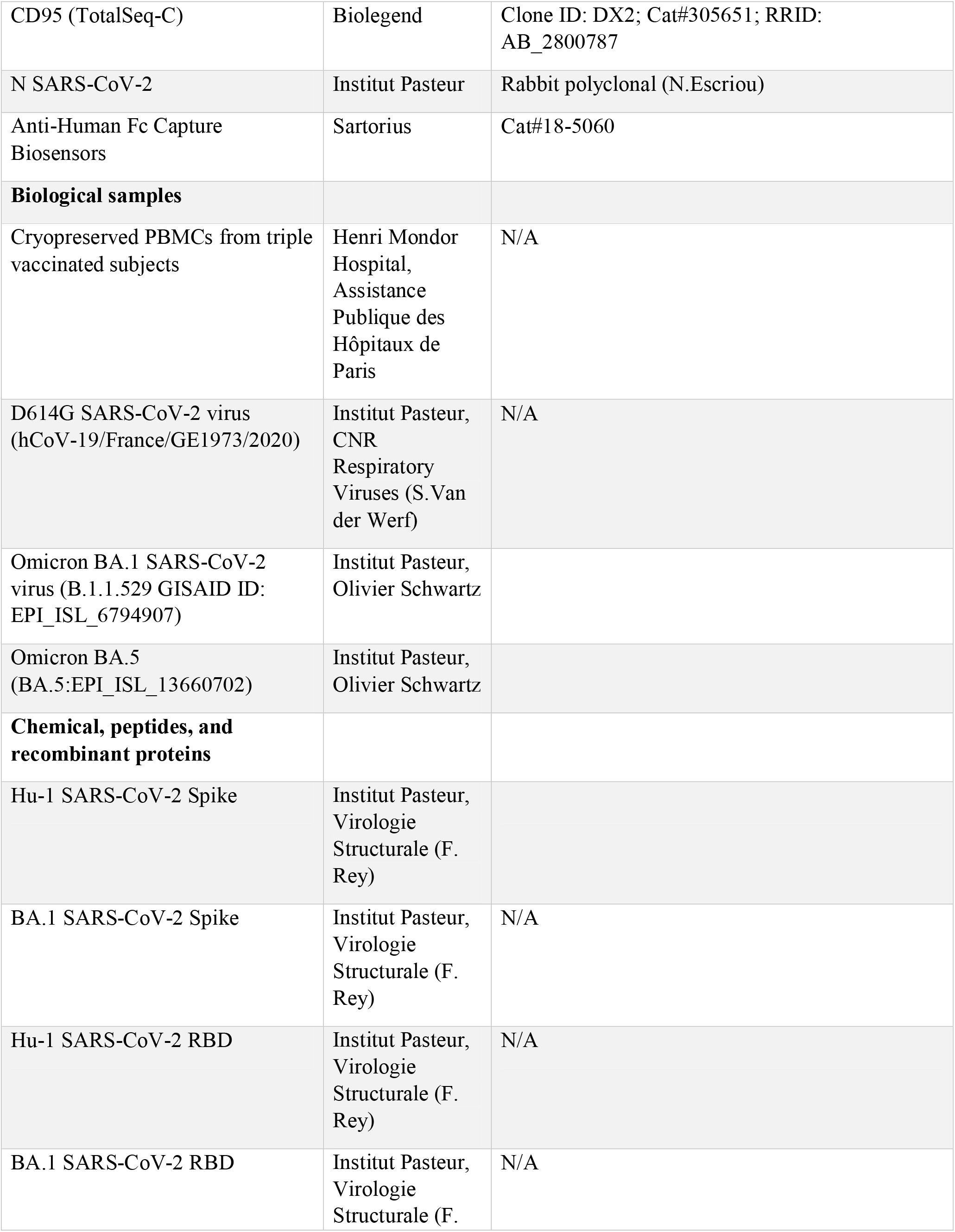
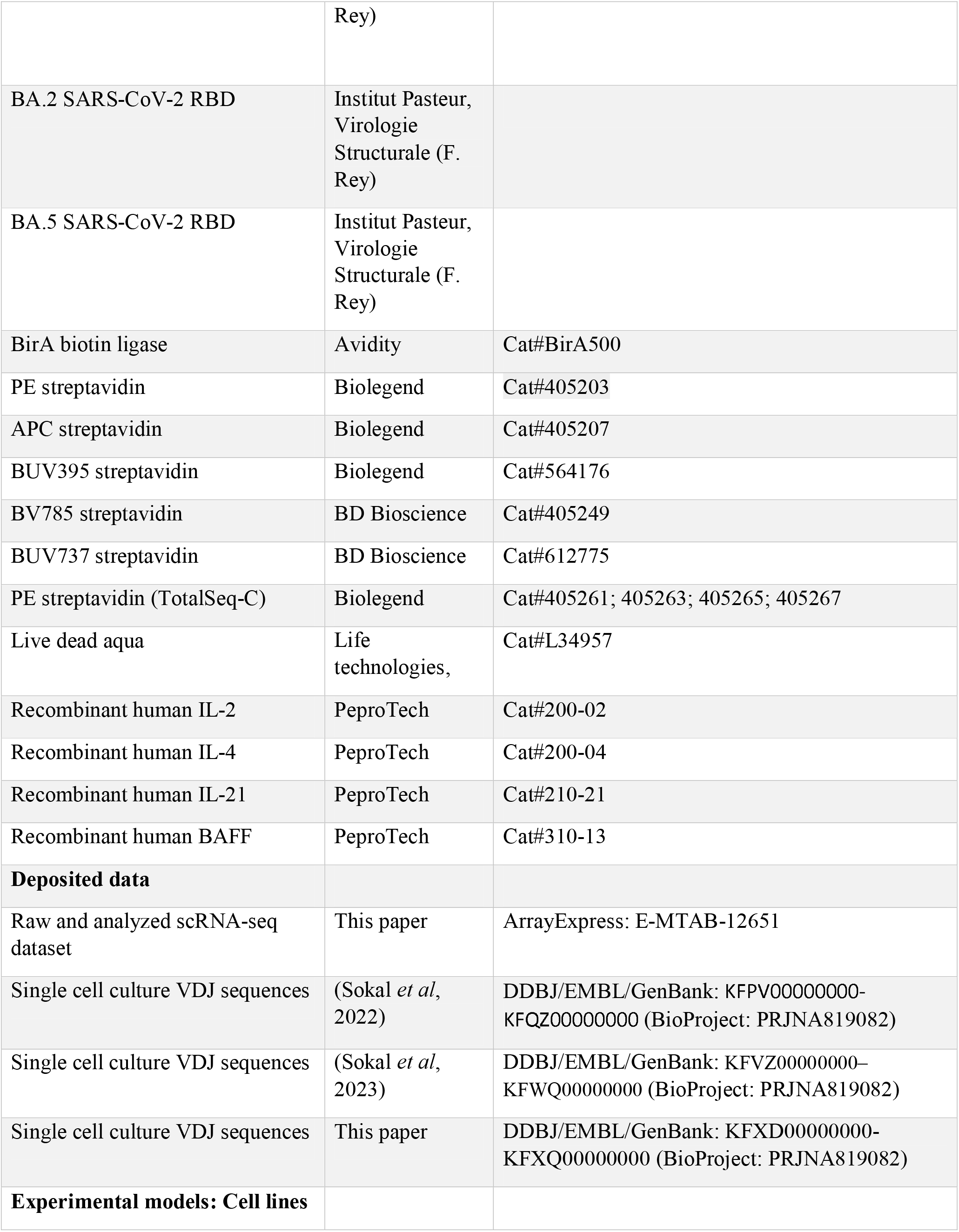
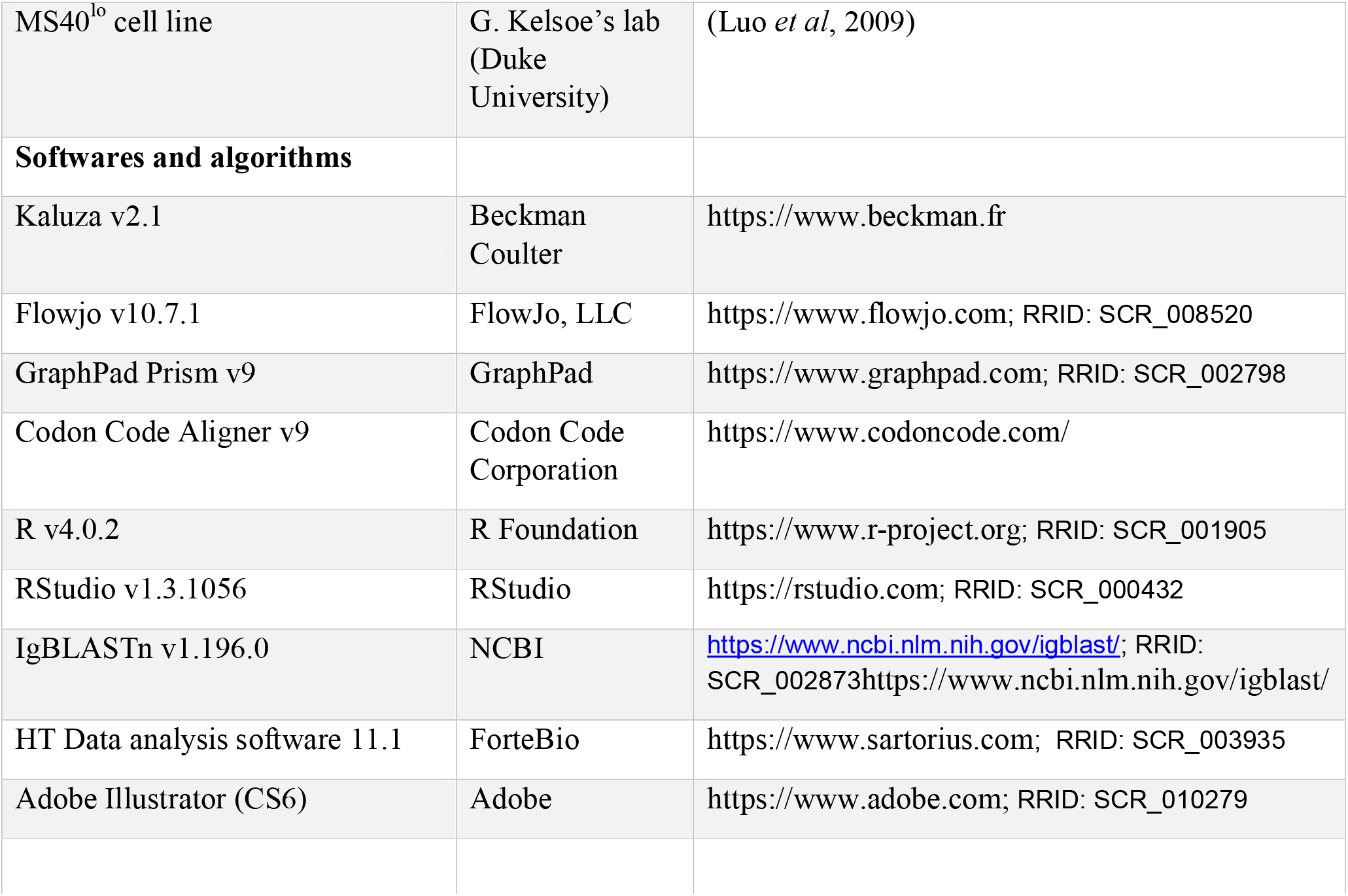

## RESOURCE AVAILABILITY

### Lead Contact

Further information and requests for resources and reagents should be directed to and will be fulfilled by the Lead Contact, Pascal Chappert.

### Materials Availability

No unique materials were generated for this study.

### Data and Code Availability

- All scRNA-seq data have been deposited in the ArrayExpress database at EMBL-EBI (www.ebi.ac.uk/arrayexpress) and will be made available as of the date of publication. Accession numbers are listed in the Key Resources Table. Single cell culture VDJ sequencing data reported in **Figure 3 and Figure S3** are included in **Table S3** and have been deposited as Targeted Locus Study projects at DDBJ/EMBL/GenBank are available as of the date of publication. Accession numbers are listed in the Key Resources Table. The version described in this paper is the first version, KFXD01000000– KFXQ01000000.
- Any additional information required to reanalyze the data reported in this paper is available from the lead contact upon request.

## EXPERIMENTAL MODEL AND SUBJECT DETAILS

### Study participants

In total, 30 patients who received a booster (3^rd^ dose) of BNT162b2 mRNA vaccine with no history of COVID-19 were enrolled. Among them, 15 developed Omicron BA.1 breakthrough infection, including 4 (Na-5, Na-9, Na-25 and Na-31) who were sampled after the third dose and before the BA.1 breakthrough infection, providing the unique opportunity of longitudinal assessment of the remodeling at the scale of one individual, especially in 2 of them whose memory B cell repertoire had been extensively characterized after 2 doses of mRNA vaccine. All the breakthrough infection occurred between 12/24/2021 and 01/30/2022 when BA.1 was responsible for > 85% of SARS-CoV-2 infections in France. They received their third dose 240 ± 40.4 (mean ± SD) days after second dose and 39 days before BA.1 breakthrough infection (range: 31 to 106 days). The remaining 15 patients were sampled at least once after their third dose of BNT162b2 mRNA vaccine, including 11 previously sampled in the MEMOCoV2 cohort (IRB 2018-A01610-55). Negative nucleocapsid IgG were assessed at each sampling during all the follow-up for these patients.

Detailed information on the individuals, including gender and health status, can be found in Table S1.

Samples were collected shortly after the boost and/or breakthrough infection (mean: 9; range: 5-12 days for 3xmRNA; mean: 14.1; range: 7-22 days for BT), 2.5 months after the boost and/or breakthrough infection (mean: 62; range: 45-79 days for 3xmRNA; mean: 75; range: 57-101 days for BT) and 5.5 months after the booster or and/or breakthrough infection (mean: 193; range: 129-227 days for 3xmRNA; mean: 164; range: 145-191 days for BT). Clinical and biological characteristics of these patients are summarized in **Table S1**.

Patients were recruited at the Henri Mondor University Hospital (AP-HP), between March 2020 and July 2022. MEMO-COV-2 study (NCT04402892) was approved by the ethical committee Ile-de-France VI (Number: 40-20 HPS) and performed in accordance with the French law. Written informed consent was obtained from all participants.

### Virus strains

The reference D614G strain (hCoV-19/France/GE1973/2020) was supplied by the National Reference Centre for Respiratory Viruses hosted by Institut Pasteur and headed by Sylvie van der Werf as described in Sokal et al. 2021. The Omicron strains (B.1.1.529 GISAID ID: EPI_ISL_6794907) and BA.5 (BA.5:EPI_ISL_13660702) were a generous gift from Olivier Schwartz, Institut Pasteur, and were generated as described in (Planas *et al*, 2021) and (Bruel *et al*, 2022) respectively.

## METHOD DETAILS

### Anti- RBD (S) and -N SARS-CoV-2 antibodies assay

Serum samples were analyzed for anti-S-RBD Hu-1 IgG titers with the SARS-CoV-2 IgG Quant II assay (ARCHITECT®, Abbott Laboratories). The latter assay is an automated chemiluminescence microparticle immunoassay (CMIA) that quantifies anti-RBD IgG, with 50 AU/mL as a positive cut-off and a maximal threshold of quantification of 40,000 AU/mL. Dilutions were performed for samples over the maximal threshold.

Serum samples were analyzed for anti-S-RBD BA.1 using the Anti-SARS-CoV-2 (B.1.1.529) Antibody IgG Titer Serologic Assay Kit (Spike RBD) kit from Acrobiosystem (RAS-T057). Sampled were diluted at 1/1000 after calibration and validation of the assay using control sera. Assay was performed according to the manufacturer instructions.

Serum samples were also processed for anti-Nucloprotein (N) detection on Abbott SARS-CoV-2 IgG chemiluminescent microparticle immunoassay following the manufacturer’s instructions.

All assays were performed by trained laboratory technicians according to the manufacturer’s standard procedures.

### Recombinant protein purification

#### Construct design

Genes coding for SARS-CoV-2 Spike (S) ectodomains (Hu-1 and BA.1) with Hisx8 and Strep tags were synthesized by Genscript and cloned into the pcDNA3.1(+) vector. Both ectodomains (residues 1-1208) were stabilized to preserve their trimeric prefusion conformation by introducing six proline substitutions (F817P, A892P, A899P, A942P, K986P, V987P, Hu-1 numbering), a GSAS substitution at the furin cleavage site (residues 682–685) and a C-terminal Foldon trimerization motif (Hsieh et al., 2020).

The SARS-CoV-2 Hu-1 and BA.1 Receptor Binding Domains (RBDs) were cloned in pcDNA3.1(+) encompassing residues 331-528 (Hu-1 numbering) from the Spike ectodomains, and they were flanked by an N-terminal IgK signal peptide and a C-terminal Thrombin cleavage site followed by Hisx8-Strep-Avi tags in tandem. The BA.2 and BA.5 RBDs were obtained using the BA.1 RBD plasmid as a template, on which the remaining mutations were introduced by PCR mutagenesis following standard techniques.

#### Protein expression and purification

The plasmids coding for the recombinant proteins were transiently transfected in Expi293F™ cells (Thermo Fischer) using FectroPRO® DNA transfection reagent (Polyplus), according to the manufacturer’s instructions. The cells were incubated at 37°C (Hu-1 S, BA.1 S, Hu-1 RBD) or 32°C (BA.1 RBD, BA.2 RBD, BA.5 RBD) for 5 days and then the culture was centrifuged, and the supernatant was concentrated. The proteins were purified from the supernatant by affinity chromatography on a StrepTactin column (IBA). The Spike proteins were further purified by size-exclusion chromatography (SEC) on a Superose6 10/300 colum (Cytiva) equilibrated in PBS, while the RBDs were loaded onto a Superdex200 10/300 column (Cytiva).

#### Protein biotinylation

Hu-1, BA.1 RBD and Spike Avi-tagged proteins were biotinylated using the Avidity BirA biotin-protein ligase kit according to the manufacturer instruction. Bovine serum albumin was biotinylated using EZ link NHS biotin (Thermofischer) according to the manufacturer instruction.

### Flow cytometry and cell sorting

PBMCs were isolated from venous blood samples via standard density gradient centrifugation and used after cryopreservation at −150°C. Cells were thawed in RPMI-1640 (Gibco)-10% FBS (Gibco), washed twice and incubated with a mixture of Hu-1 and BA.1 Spike +/− Hu-1 and BA.1 RBD tetramers in 100 μL of PBS (Gibco)-2% FBS during 40 min on ice. For cell sorting, cells were stained with 500 ng of Hu-1Spike APC-streptavidin and 500 ng of BA.1 Spike PE-streptavidin; for flow cytometry analysis cell were stained with 500 ng of Hu-1Spike BUV395-streptavidin and 500 ng BA.1 Spike PE-streptavidin 50ng of Hu-1 RBD APC-streptavidin and 50 ng of BA.1 RBD BV785 Streptavidin. To exclude cells with nonspecific binding, a non-relevant tetramer was constructed using biotinylated bovine serum albumin coupled to BV785-streptavidin (for cell sorting) or BU737-streptavidin for flow cytometry. Tetramer were made by incubating biotinylated proteins with fluorochrome-conjugated streptavidin at 4:1 molar ratio for 1 hour at 4°C. 2.4 ng of free biotin was then added for 10 additional minutes before mixing of the tetramer. Cells were then washed and resuspended in the same conditions, then the fluorochrome-conjugated antibody cocktail at pre-titrated concentrations (1:100 for CD19, CD21, CD11c, CD71, CD38, CD3, CD14 and IgD, 1:50 for CD27) for 20 min at 4°C and viable cells were identified using a LIVE/DEAD Fixable Aqua Dead Cell Stain Kit (Thermo Fisher Scientific, 1:200) incubated with conjugated antibodies. Samples were acquired using a LSR Fortessa SORP (BD Biosciences). For cell sorting, cells were stained using the same protocol and then sorted in 96 plates using the ultra-purity mode on a MA900 (SONY) or an Aria II cell sorter (BD Biosciences) and Data were analyzed using FlowJo or Kaluza softwares. Detailed gating strategies for cell sorting and analysis are depicted in **Figures S1 and S2** respectively and in **Table S3B**.

For UMAP generation and visualization (**Figure 2 and Figure S2**), viable dump^−^ CD19^+^ IgD^−^ cells from each sample included in the final analysis (**Table S1**) were first downsampled to 4000. The UMAP (v3.1) plugin in FlowJO was then used on a concatenated FCS file containing all donors and time points to calculate the UMAP coordinates for the resulting 264.000 cells (with 30 neighbors, metric = euclidian and minimum distance = 0.5 as default parameters), considering fluorescent intensities from the following parameters: FSC-A, SSC-A, CD19, CD21, CD11c, CD71, CD38, CD27 and IgD, while excluding the dump (CD3 and CD14), viability and Tetramers channels. Contour plots (equal probability contouring, with intervals set to 5% of gated populations) for each manually gated populations (**Figure S2A**) were further overlaid on UMAP projection in FlowJO (**Figure S2B**). For visualization purposes, only the outermost density representing 95% of the total gated cells was kept for the final figure, all other contour lines were removed in Adobe Illustrator.

### Single-cell culture

Single cell culture was performed as previously described (Crickx et al., 2021). Single B cells were sorted in 96-well plates containing MS40L^lo^ cells expressing CD40L (kind gift from G. Kelsoe, Luo et al., 2009). Cells were co-cultured at 37°C with 5% CO2 during 21 or 25 days in RPMI-1640 (Invitrogen) supplemented with 10% HyClone FBS (Thermo Scientific), 55 μM 2-mercaptoethanol, 10 mM HEPES, 1 mM sodium pyruvate, 100 units/mL penicillin, 100 μg/mL streptomycin, and MEM non-essential amino acids (all Invitrogen), with the addition of recombinant human BAFF (10 ng/ml), IL2 (50 ng/ml), IL4 (10 ng/ml), and IL21 (10 ng/ml; all Peprotech). Part of the supernatant was carefully removed at days 4, 8, 12, 15 and 18 and the same amount of fresh medium with cytokines was added to the cultures. After 25 days of single cell culture, supernatants were harvested and stored at −20°C. Cell pellets were placed on ice and gently washed with PBS (Gibco) before being resuspended in 50 μL of RLT buffer (Qiagen) supplemented with 1% β-mercaptoethanol and subsequently stored at −80°C until further processing.

### ELISA

Total IgG and SARS-CoV-2 Hu-1 RBD, Hu-1 S, BA.1 RBD and BA.1 S-specific IgG from culture supernatants were measured using homemade ELISA. 96 well ELISA plates (Thermo Fisher) were coated with either goat anti-human Ig (10 μg/ml, Invitrogen) or recombinant SARS-CoV-2 Hu-1 −RBD or −S or BA.1-RBD or −S protein (2.5 μg/ml each) in sodium carbonate during 1h at 37°C. After plate blocking, cell culture supernatants were added for 1hr, then ELISA were developed using HRP-goat anti-human IgG (1 μg/ml, Immunotech) and TMB substrate (Eurobio). OD450 and OD620 were measured, and Ab-reactivity was calculated after subtraction of blank wells. Supernatants whose ratio of OD450-OD620 over control wells (consisting of supernatant from wells that contained spike-negative MBCs from the same single cell culture assay) was over 10 were considered as positive for Hu-1 RBD or BA.1 RBD. PBS was used to define background OD450-OD620.

### Single-cell RNA-seq library preparation and sequencing

Frozen PBMC from 4 donors (Na-9, Na-25, Na-37 and Na-38) were thawed and washed 2 times as described above. 10-15×10^6^ PMBCs were then resuspended in 100μL PBS 2%FBS and incubated for 40 minutes at 4°C with a decoy tetramer (biotinylated Bovine Serum albumin coupled with BV785 streptavidin) and Hu-1 Spike, BA.1 Spike, Hu-1 RBD and BA.1 RBD tetramers (constructed as described above using PE-labelled Total-seqC streptavidin with different barcodes for each individual antigens). Cell were washed, resuspended in 100μL PBS 2%FBS and stained with a cocktail of fluorochrome conjugated (CD3, CD14 both APC-H7 at 1:100 each; CD15 and CD56 BV785 at 1:100 each, CD19 PECF594 at 1:100, IgD FITC at 1:100, CD38 PercP-Cy5.5 at 1:100) and CITE-seq (CD38, CD27, CD71, CD21, CD11c, CD39, FCRL5, CD95 all at 1:40) antibodies for 40 minutes on ice. Viable cells were identified using a LIVE/DEAD Fixable Aqua Dead Cell Stain Kit (Thermo Fisher Scientific, 1:200) incubated with conjugated antibodies. B cells were FACS-sorted (MA900, Sony) in PBS/0.08% FCS from 4 patients at baseline (M0) and 6 months (M6). An initial pool of 50.000 total CD19^+^IgD^−^ cells were always sorted and afterward, to enrich for cells of interest, only CD19^+^CD38^low^ antibody secreting cells (ASCs), PE/tetramer positive and CD19^hi^ cells, leading to approximately 55000-60000 total sorted cells per sample. Sorted cells were then counted and up to 20 000 cells were loaded in the 10x Chromium Controller to generate single-cell gel-beads in emulsion. The scRNA-seq libraries for gene expression (mRNA), ADT and VDJ BCR libraries were generated using Chromium Next GEM Single Cell V(D)J Reagent Kit v.1.1 with Feature Barcoding (10x Genomics) according to the manufacturer’s protocol. Briefly, after reverse transcription, gel-beads in emulsion were disrupted. Barcoded complementarity DNA was isolated and amplified by PCR. Following fragmentation, end repair and A-tailing, sample indexes were added during index PCR. The purified libraries were sequenced on a Novaseq S2 flowcell (Illumina) with 26 cycles of read 1, 8 cycles of i7 index and 91 cycles of read 2, targeting a median depth of 50000 reads per cell for gene expression and 5000 reads per cell for each other two libraries (BCR VDJ and ADT Feature barcoding).

### Single-cell IgH sequencing

Clones whose culture had proven successful (IgG concentration ≥ 1 μg/mL at day 21-25) were selected and extracted using the NucleoSpin96 RNA extraction kit (Macherey-Nagel) according to the manufacturer’s instructions. A reverse transcription step was then performed using the SuperScript IV enzyme (ThermoFisher) in a 14 μl final volume (42°C 10 min, 25°C 10 min, 50°C 60 min, 94°C 5 min) with 4 μl of RNA and random hexamers (Thermofisher scientific). A PCR was further performed based on the protocol established by Tiller et al (Tiller *et al*, 2008). Briefly, 3.5 μl of cDNA was used as template and amplified in a total volume of 40 μl with a mix of forward L-VH primers (**Table S3**) and reverse Cγ primer and using the HotStar® Taq DNA polymerase (Qiagen) and 50 cycles of PCR (94°C 30 s, 58°C 30 s, 72°C 60 s). PCR products were sequenced with the reverse primer CHG-D1 and read on ABI PRISM 3130XL genetic analyzer (Applied Biosystems). Sequence quality was verified using CodonCode Aligner software (CodonCode Corporation).

For specific patients and time points (see **Table S1**), some IgH sequences were obtained directly from single cell sorting in 4μL lysis buffer containing PBS (Gibco), DTT (ThermoFisher) and RNAsin (Promega). Reverse transcription and a first PCR was performed as described above (50 cycles) before a second 50-cycles PCR using 5’AgeI VH primer mix and Cγ-CH1 3’ primer, before sequencing.

### Single-cell gene expression analysis

Paired end FASTQ reads for all three libraries were demultiplexed and aligned against the GRCh38 human reference genome (GENCODE v32/Ensembl 98; July 2020) using 10x Genomics’ Cell Ranger v6.0.0 pipeline. Outputs of Cell Ranger were directly loaded into Seurat v4.1.1(Hao *et al*, 2021) for further QC steps and analysis. Following manual inspection of cell quality, only genes detected in at least 10 cells and cells with more than 750 unique genes detected and less than 5% of UMI counts mapped to mitochondrial genes were kept (**Figure S1B)**. Upon analysis of parallel VDJ library (see “Computational analyses of VDJ sequences” section below), only cells with exactly one resolved heavy chain sequence were retained for final analysis. Transcript counts were first normalized using the scTransform algorithm v0.3.4 (Hafemeister & Satija, 2019), using the vst.flavor “v2” parameter and additionally correcting for potential bias related to the detected percentage of mitochondrial genes and selecting for the top 3000 variable features for downstream visualization and clustering analysis. After principal component analysis, performed excluding all remaining Ig genes to avoid unwanted clustering based solely on differential isotype expression (https://www.genenames.org/data/genegroup/#!/group/348 and AC233755.1 gene), potential donor and sort-specific batch effects were removed using the Harmony algorithm (Korsunsky *et al*, 2019). The first 15 corrected PCA dimensions were then used to construct a knn graph (k=20 neighbors) and perform graph-based clustering (Louvain) with a resolution parameter of 0.2 as well as compute the UMAP coordinates for each cell. G2M and S cell cycle signatures were calculated using the CellCycleScoring() function and the associated gene lists in Seurat (G2M scoring: HMGB2, CDK1, NUSAP1, UBE2C, BIRC5, TPX2, TOP2A, NDC80, CKS2, NUF2, CKS1B, MKI67, TMPO, CENPF, TACC3, FAM64A, SMC4, CCNB2, CKAP2L, CKAP2, AURKB, BUB1, KIF11, ANP32E, TUBB4B, GTSE1, KIF20B, HJURP, CDCA3, HN1, CDC20, TTK, CDC25C, KIF2C, RANGAP1, NCAPD2, DLGAP5, CDCA2, CDCA8, ECT2, KIF23, HMMR, AURKA, PSRC1, ANLN, LBR, CKAP5, CENPE, CTCF, NEK2, G2E3, GAS2L3, CBX5, CENPA; S scoring: MCM5, PCNA, TYMS, FEN1, MCM2, MCM4, RRM1, UNG, GINS2, MCM6, CDCA7, DTL, PRIM1, UHRF1, MLF1IP, HELLS, RFC2, RPA2, NASP, RAD51AP1, GMNN, WDR76, SLBP, CCNE2, UBR7, POLD3, MSH2, ATAD2, RAD51, RRM2, CDC45, CDC6, EXO1, TIPIN, DSCC1, BLM, CASP8AP2, USP1, CLSPN, POLA1, CHAF1B, BRIP1, E2F8).

### Computational analyses of VDJ sequences

Processed FASTA sequences from cultured single-cell heavy chain sequencing and 10x single-cell RNA sequencing were annotated using Igblast v1.19.0 against the human IMGT reference database (**Figure S1B**). Sequences from cells that did not pass the initial QC cutoffs from our scRNA-seq analysis were removed at that step. Cases of 10x barcodes with two or more consensus heavy chain sequences for which more than ten UMI were detected were generally flagged as potential doublets for removal from our scRNA-seq analysis. Similarly, cases where no clear heavy chains could be attributed (none above 10 UMIs) were also flagged for removal. Two exceptions were made: 1/ in cases of identical CDR3s but differing isotypes (c_call), in which case the isotype switched sequence was kept and UMI counts from both contigs were aggregated; and 2/ in cases when one the heavy chains was clearly overrepresented at the UMI level (at least three time the number of UMI counts as compared to the next most represented heavy chain detected) and the second most represented sequences did not exceed ten UMIs, in which case the most represented sequence was kept.

Clonal cluster assignment (DefineClones.py) and germline reconstruction (CreateGermlines.py) was performed using the Immcantation/Change-O toolkit (Gupta *et al*, 2015) on all heavy chain V sequences. Sequences that had the same V-gene, same J-gene, including ambiguous assignments, and same CDR3 length with maximal length normalized nucleotide hamming distance of 0.15 were considered as potentially belonging to the same clonal group. Mutation frequencies in V genes were then calculated using the calcObservedMutations() function from Immcantation/SHazaM v1.1.1 R package. For the analysis of the initial ASC response in our 10x dataset (**Figure 2E/F**), clonal assignments were further corrected using available light chain information (light_cluster.py script from Immcantation). Further clonal analyses on all productively rearranged sequences were implemented in R.

Based on heavy-chain only clonal affectation, clones were defined as Hu-1 or BA.1 SARS-Cov-2 S or RBD-specific if they contained 1 or more validated single-cell culture sequence or cells positively stained by our barcoded His-tagged S or RBD protein in our scRNAseq dataset. Staining with barcoded S or RBD tetramer in our scRNAseq dataset were analyzed in FlowJO using log-normalized sequencing data (see **Figure S1B**). Clones containing RBD-specific cells were labelled as RBD-specific. Clones containing Hu-1/BA.1 cross-reactive cells were labelled as cross-reactive. All specific clones were manually curated based on available light chain information and CDR3 sequences and clones containing less than ten percent of barcoded S or RBD tetramer-stained cells and no in vitro validated cells were manually labeled as unknown specificity.

Clones from which members were found before and after BA.1 BT infection were labelled as “sustained”. Clones seen at least twice before BA.1 BT infection, never after BA.1 BT infection and whose overall frequency pre-BA.1 BT infection was superior to the frequency of singletons post-BA.1 BT infection in that donor, to account for differences in sampling pre-and post- BA.1 BT infection, were labelled as “lost”. Clones never seen before BA.1 BT infection, seen at least twice after BA.1 BT infection and whose overall frequency post-BA.1 BT infection was superior to the frequency of singletons pre-BA.1 BT infection in that donor were labelled as “newly expanded”. Both “lost” and “newly expanded” clones were further labelled as “persisting” if found at multiple time points.

VH repartitions and Shannon entropies were calculated using the countGenes() and alphaDiversity() functions from the Immcantation/alakazam v1.2.0 R package. Chao1 richness indexes were calculated using the iNEXT v3.0.0 package (https://doi.org/10.1111/2041-210X.12613). Identical sequences were identified using the collapseDuplicates() function from the Immcantation/alakazam v1.2.0 R package. To account for differences in sampling, we computed the average frequency of duplicates found upon 1000 bootstrapping, downsampling to 65 sequences per time point for each donor.

Phylogenetic trees and date randomization test to detect evolution over time were generated and performed using the Dowser v1.0.0 package (Hoehn *et al*, 2022) and the Immcantation/IgPhyML toolkit (Immcantation/suite v4.3.2) and further visualized in R using the Alakazam v1.2.0 and ggtree v3.4.2 packages.

Graphics were obtained using the ggplot2 v3.3.6, pheatmap v1.0.12 and circlize v0.4.15 packages.

### Affinity measurement using biolayer interferometry (Octet)

This high-throughput kinetic screening of supernatants using single antigen concentration has recently been extensively tested and demonstrated excellent correlation with multiple antigen concentration measurements (Lad et al., 2015). Biolayer interferometry assays were performed using the Octet HTX instrument (ForteBio). Anti-Human Fc Capture (AHC) biosensors (18-5060) were immersed in supernatants from single-cell MBC cultures (or control monoclonal antibody) at 25°C for 1800 seconds. Biosensors were equilibrated for 10 minutes in 10x PBS buffer with surfactant Tween 20 (Xantec B PBST10-500) diluted 1x in sterile water with 0.1% BSA added (PBS-BT) prior to measurement. Association was performed for 600s in PBS-BT with Hu-1 or variant RBD (BA.1, BA.2 and BA.5) at 100nM followed by dissociation for 600s in PBS-BT. Biosensor regeneration was performed by alternating 30s cycles of regeneration buffer (glycine HCl, 10 mM, pH 2.0) and 30s of PBS-BT for 3 cycles. Traces were reference sensor subtracted and curve fitting was performed using a local 1:1 binding model in the HT Data analysis software 11.1 (ForteBio). Sensors with response values (maximum RBD association) below 0.1nm were considered non-binding. Hu-1 RBD non-binding monoclonal antibodies (n=14/414) were excluded from further analysis. For variant RBD non-binding mAbs, sensor-associated data (mAb loading and response) were manually checked to ensure that this was not the result of poor mAb loading. For binding clones, only those with full R^2^>0.8 were retained for KD reporting and Omicron lineage binding. mAbs were defined as affected against a given variant RBD if the ratio of calculated KD value against that RBD variant and the Hu-1 RBD was superior to five and final KD > 5×10^−10^ M. Omicron lineage binding residues prediction was simply made based on mutations repartition in the different variants.

### Virus neutralization assay

Virus neutralization was evaluated by a focus reduction neutralization test (FRNT). Vero E6 cells were seeded at 2×10^4^ cells/well in a 96-well plate 24h before the assay. Two-hundred focus-forming units (ffu) of virus were pre-incubated with serial dilutions of heat-inactivated sera for 1hr at 37°C before infection of cells for 2hrs or with supernatants from single-cell cultured memory B cells at 16nM. The virus/antibody mix was then removed, and foci were left to develop in presence of 1.5% methylcellulose for 2 days (D614G) or 3 days (Omicron BA.1 and BA.5). Cells were fixed with 4% formaldehyde and foci were revealed using a rabbit anti-SARS-CoV-2 N antibody (gift of Nicolas Escriou) and anti-rabbit secondary HRP-conjugated secondary antibody. Foci were visualized by diaminobenzidine (DAB) staining and counted using an Immunospot S6 Analyser (Cellular Technology Limited CTL). Pre-pandemic serum (March 2012) was used as negative control for sera titration and was obtained from an anonymous donor through the ICAReB platform (BRIF code n°BB-0033-00062) of Institut Pasteur that collects and manages bioresources following ISO (International Organization for Standardization) 9001 and NF S 96-900 quality standards.

Percentage of virus neutralization was calculated as (100 − ((#foci sample / #foci control)*100)). Sera IC50 were calculated over 8 four-fold serial dilutions from 1/10 to 1/164000 using the equation log (inhibitor) vs. normalized response – Variable slope in Prism 9 (GraphPad software LLC).

### Quantification and Statistical Analysis

Ordinary One-way ANOVA, Two-way ANOVA, Repeated measures mixed effects model analysis, Kruskal-Wallis test and Mann-Whitney test were used to compare continuous variables as appropriate (indicated in Figures). Benjamini, Krieger and Yekutieli FDR correction was used for all multiple comparisons. A *P*-value ≤ 0.05 was considered statistically significant. Statistical analyses were all performed using GraphPad Prism 9.0 (La Jolla, CA, USA).

### Additional resources

ClinicalTrials.gov Identifier: MEMO-CoV2, NCT04402892.

### Excel table titles

Table S1. Human donor’s information, experimental inclusion and serological data, related to all Figures and supplementary Figures.

Table S2. Flow cytomety-related data. Related to Figures 2 and Figures S2.

Table S3. RBD and S-specific memory B cell VDJ sequences and ELISA, affinity and neutralization data. Related to Figures 2, 3, 4, 5, 6 and Figures S2, S3, S4, S5, S6.

