## Supplementary figures and images for "Omicron BA.1 breakthrough infection drives long-term remodeling of the memory B cell repertoire in vaccinated individuals"

### Figure S1

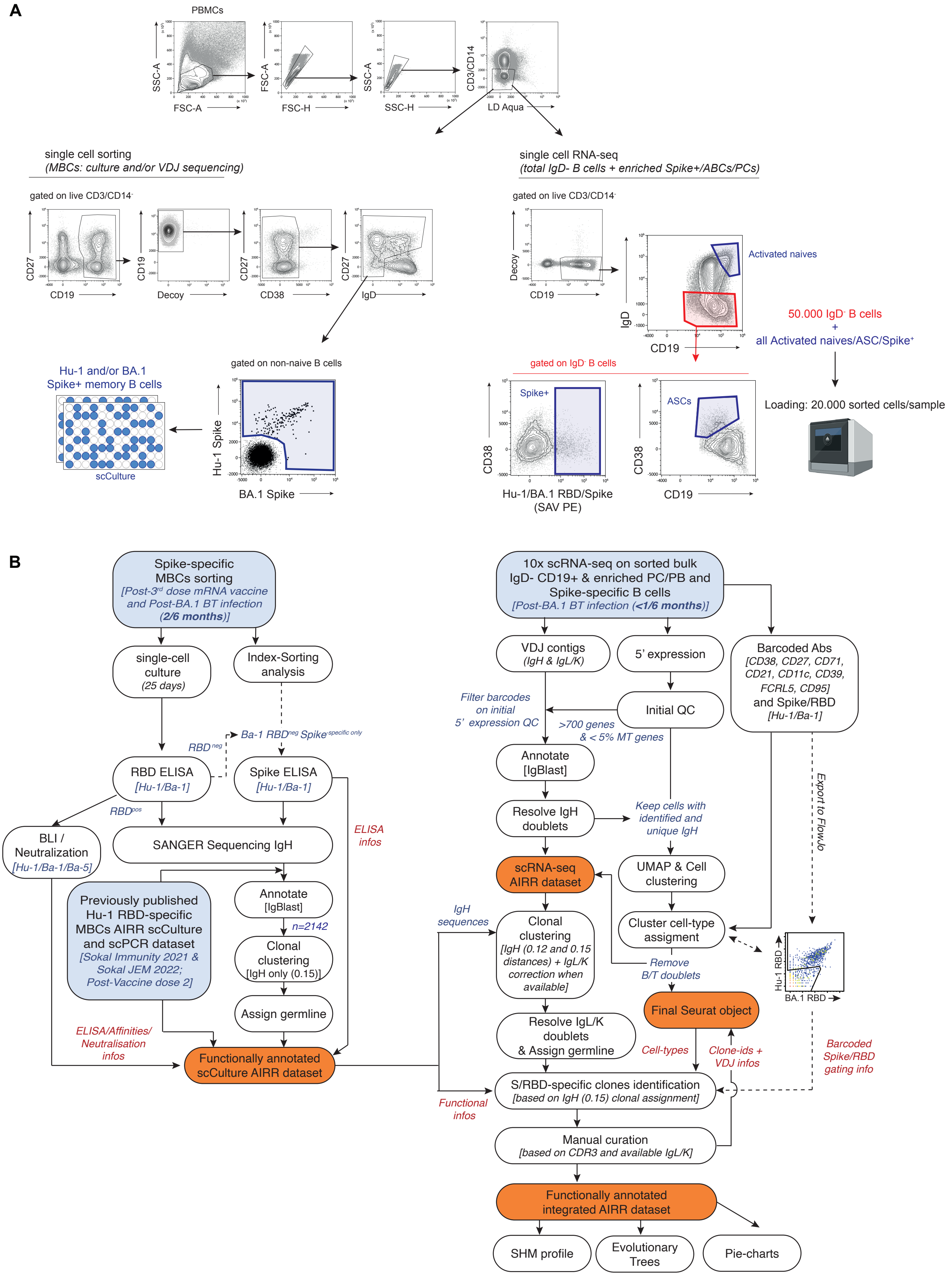

### Figure S2

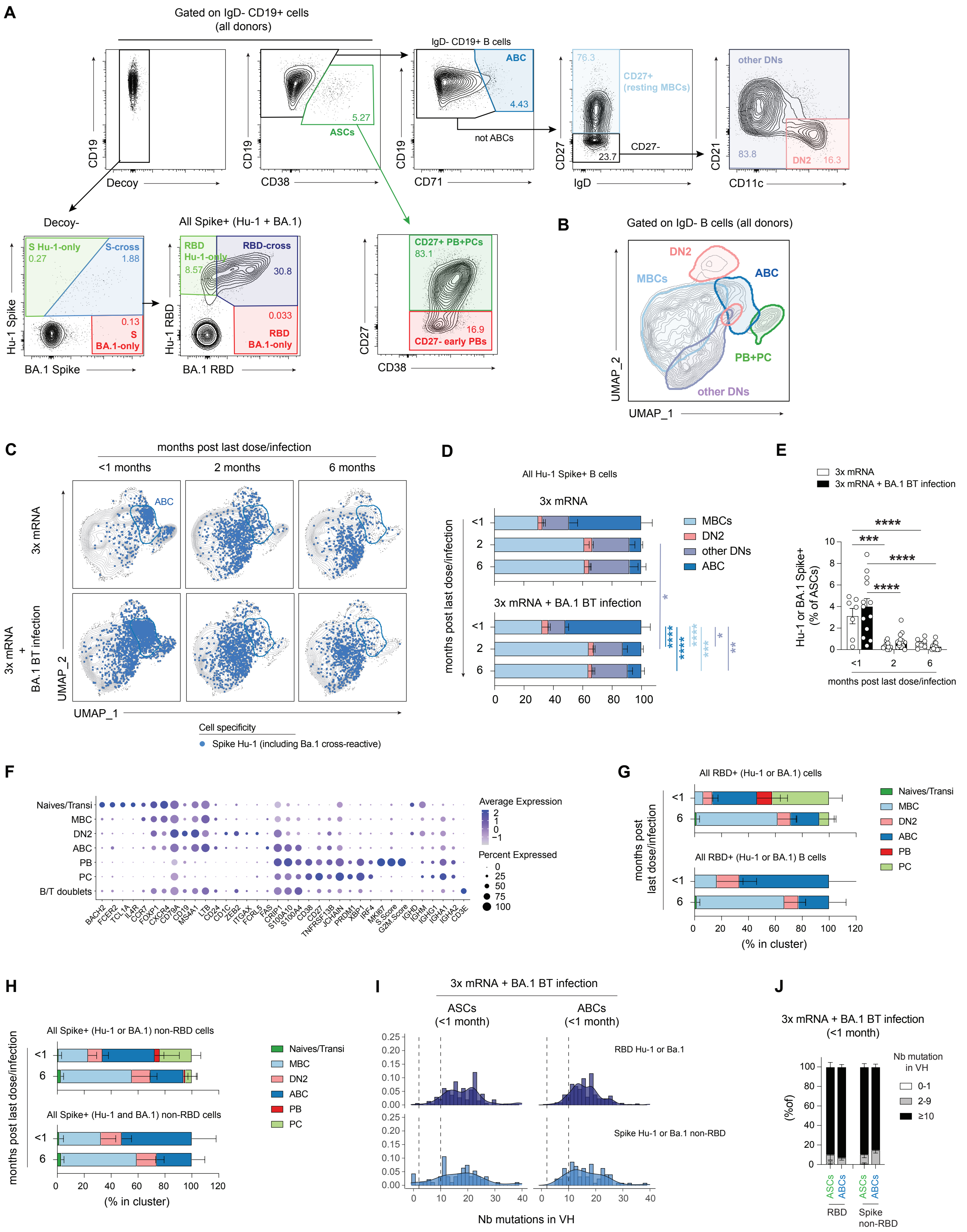

### Figure S3

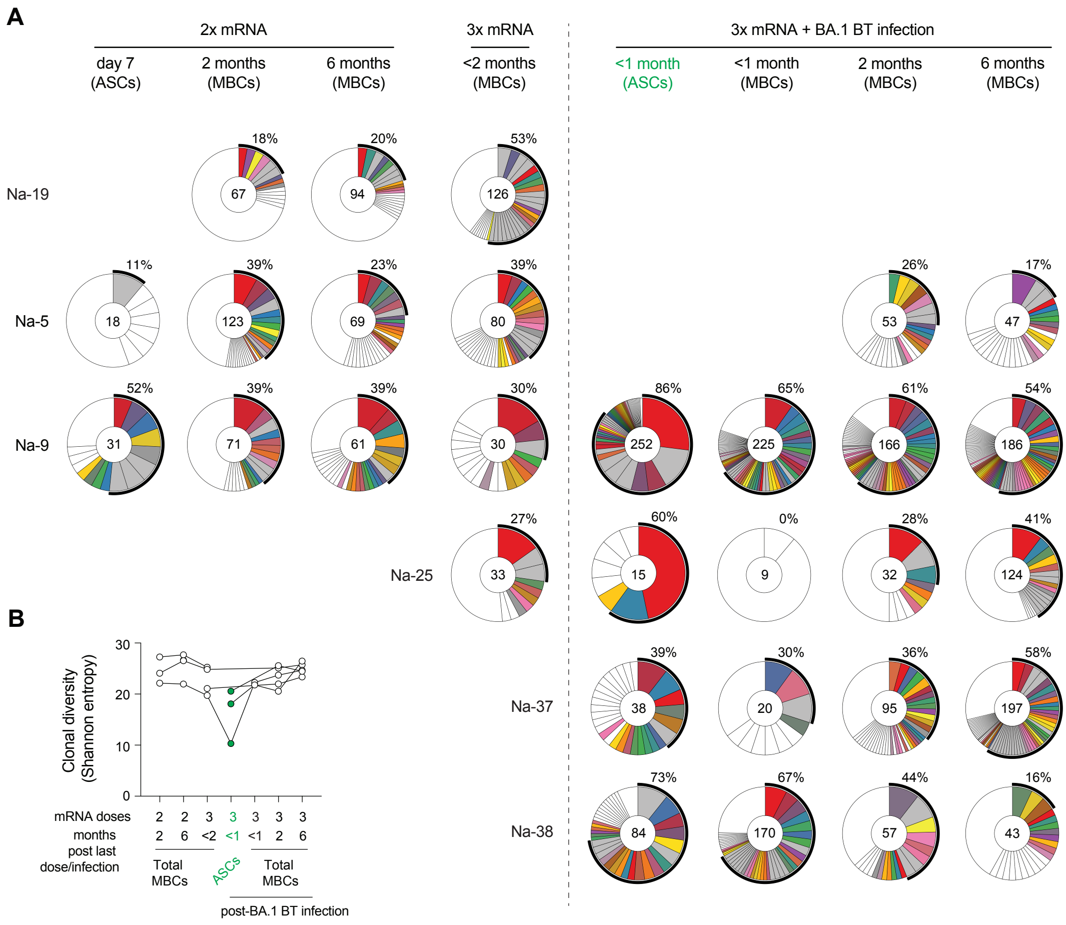

### Figure S5

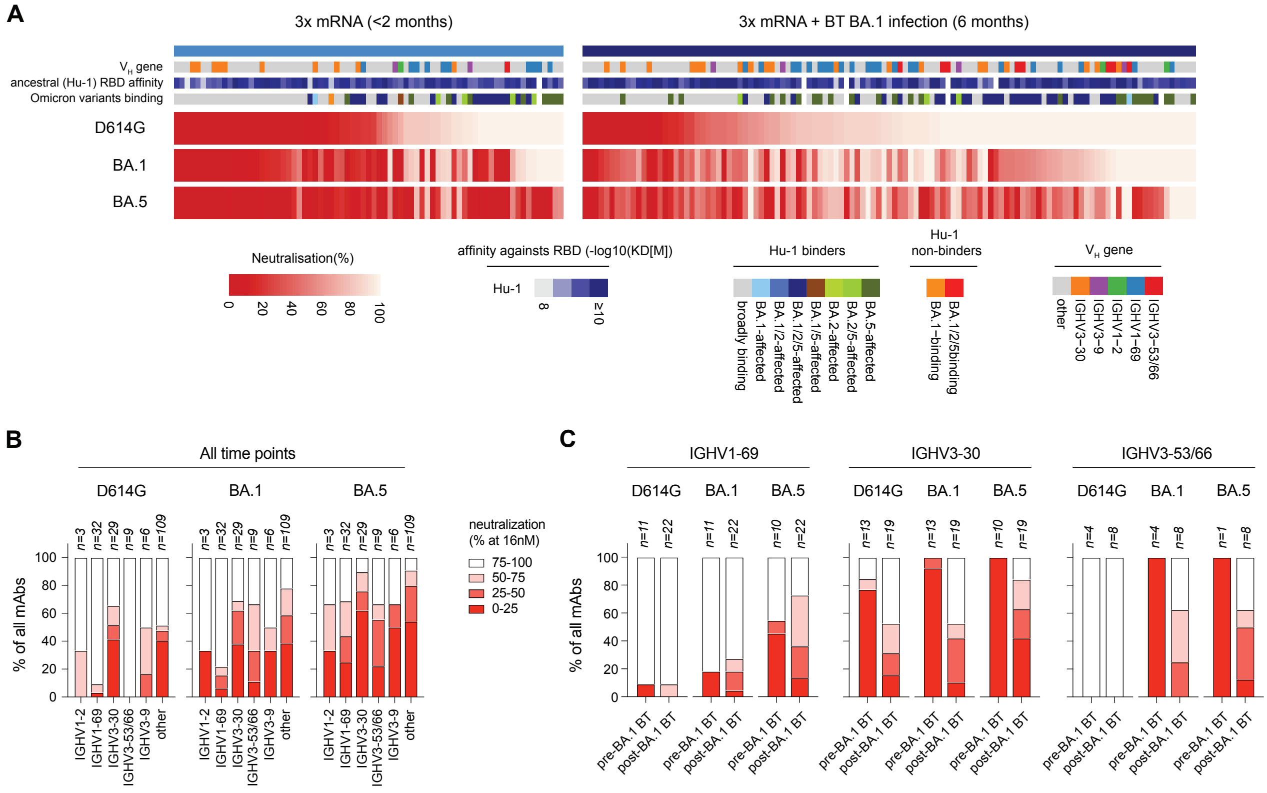

### Figure S6

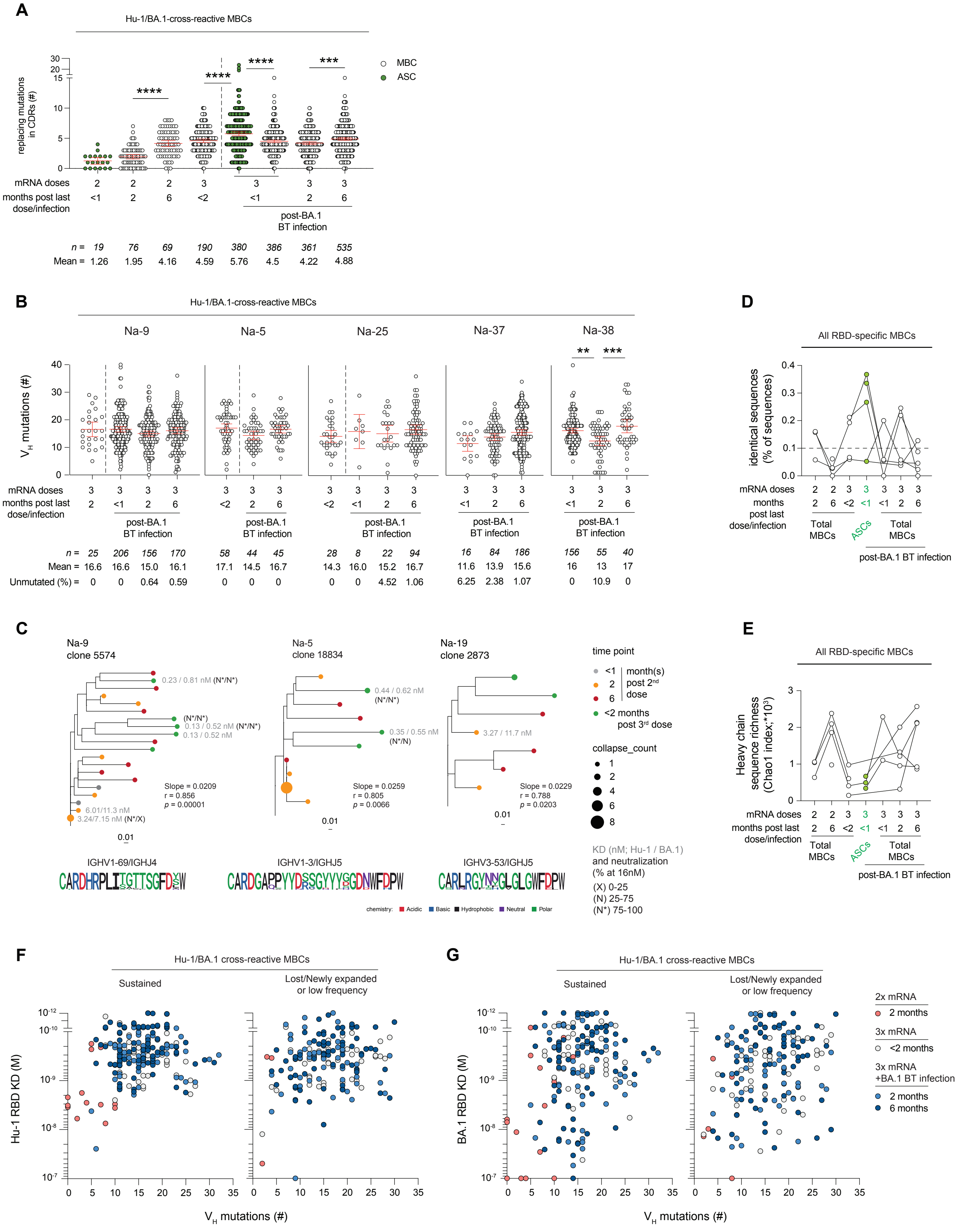

### Fiugre S4

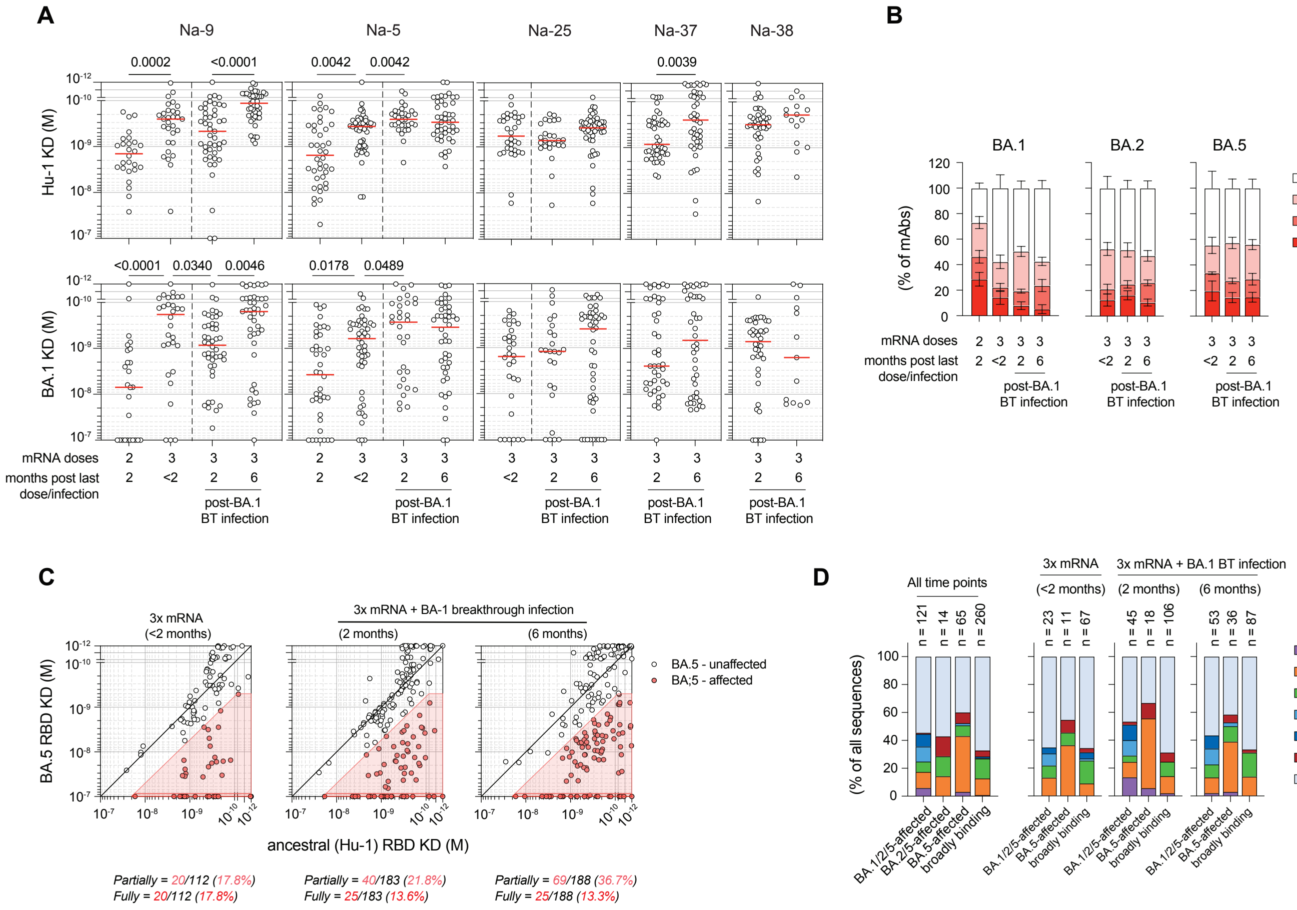
